# A Reference Atlas of the Human Dorsal Root Ganglion

**DOI:** 10.1101/2025.11.05.686654

**Authors:** Shamsuddin A. Bhuiyan, Saad S. Nagi, Ishwarya Sankaranarayanan, Evangelia Semizoglou, Dmitry Usoskin, Lite Yang, Huasheng Yu, Asta Arendt-Tranholm, Zachariah Bertels, Parth Bhatia, Otmane Bouchatta, Kevin Boyer, Anna Cervantes, Joshua Chalif, Himanshu Chintalapudi, Anthony Cicalo, Bryan Copits, Caitlin Cronin, Michele Curatolo, Xianjun Dong, Patrick M. Dougherty, Adam Dourson, Geoffrey Funk, Katherin Gabriel, Dustin S. Griesemer, Hong Guo, Prashant Gupta, Christoph Hofstetter, Peter Horton, Amanda Hsieh, Nikhil Nageshwar Inturi, Aakanksha Jain, Selwyn Jayakar, Benjamin Johnston, Rebecca Kim, Doris Krauter, Jussi Kupari, John Lemen, Joseph B. Lesnak, Weiqiang Liu, Iris Lopez, Yi Lu, Hannah J. MacMillan, Khadijah Mazhar, Pauline Meriau, Jeffrey R. Moffitt, Marisol Mancilla Moreno, Juliet M. Mwirigi, Huma Naz, Jayden O’Brein, Maria Payne, John Del Rosario, Sarah F. Rosen, Stephanie Shiers, Ebenezer Simpson, Richard Slivicki, James R. Stone, Diana Tavares-Ferreira, Megan Uhelski, Clifford J. Woolf, Qingru Xu, Jiwon Yi, Muhammad Saad Yousuf, Difei Zhu, Valeria Cavalli, Guoyan Zhao, Håkan Olausson, Patrik Ernfors, Robert W. Gereau, Wenqin Luo, Theodore J. Price, William Renthal, NIH PRECISION Human Pain Network

**Affiliations:** Department of Neurology, Brigham and Women’s Hospital and Harvard Medical School. Boston, MA, USA; Department of Biomedical and Clinical Sciences, Linköping University, Linköping, Sweden; Department of Neuroscience and Center for Advanced Pain Studies, University of Texas at Dallas, Richardson, TX, USA; Department of Medical Biochemistry and Biophysics, Division of Molecular Neurobiology, Karolinska Institute, Stockholm, Sweden; Washington University Pain Center and Department of Anesthesiology, Washington University School of Medicine, St Louis, Missouri 63110, USA; Department of Neuroscience, Perelman School of Medicine, University of Pennsylvania, Philadelphia, PA, USA; Department of Genetics, Washington University School of Medicine, St Louis, Missouri 63110, USA; Southwest Transplant Alliance, Dallas, TX, USA; Department of Neurosurgery, Brigham and Women’s Hospital and Harvard Medical School. Boston, MA, USA; Department of Neurology, Yale School of Medicine, Yale University, 100 College St, New Haven, CT, USA; Department of Anesthesiology and Pain Medicine, University of Washington, Seattle, WA, USA; Department of Pain Medicine, University of Texas MD Anderson Cancer Center, Houston, TX, USA; Department of Anesthesiology, Brigham and Women’s Hospital and Harvard Medical School. Boston, MA, USA; F.M. Kirby Neurobiology Center and Department of Neurobiology, Boston Children’s Hospital and Harvard Medical School, Boston, MA 02115, USA; Department of Neuroscience, Washington University School of Medicine, St. Louis, MO, USA; Program in Cellular and Molecular Medicine and Department of Microbiology, Boston Children’s Hospital and Harvard Medical School, Boston, MA, USA; Department of Microbiology, Blavatnik Institute, Harvard Medical School, Boston, MA, USA; Department of Pathology, Massachusetts General Hospital and Harvard Medical School, Boston, MA, USA; Department of Genetics and Department of Neurology, Hope Center for Neurological Disorders, Washington University School of Medicine, St Louis, Missouri 63110, USA

**Author notes:** equal contribution listed alphabetically (Shamsuddin A. Bhuiyan, Saad S. Nagi, Ishwarya Sankaranarayanan, Evangelia Semizoglou, Dmitry Usoskin, Lite Yang, Huasheng Yu). senior authors (Valeria Cavalli, Guoyan Zhao, Håkan Olausson, Patrik Ernfors, Robert W. Gereau, Wenqin Luo, Theodore J. Price, William Renthal).

## Abstract

Somatosensory perception largely emerges from diverse peripheral sensory neurons whose cell bodies reside in dorsal root ganglia (DRG). Damage or dysfunction of DRG neurons is a major cause of chronic pain and sensory loss. In mice, deep single-cell transcriptomic profiling and genetically defined models have offered important clues into DRG function, but in humans, the cellular and molecular landscape of DRG neurons remains less understood. Here, we constructed a reference cell atlas of the human DRG by profiling transcriptomes of cells and nuclei from 126 donors sampled across cervical, thoracic, and lumbar DRGs. This atlas resolves 22 neuronal subtypes, including known and previously unrecognized subtypes linked to nociception, mechanosensation, thermosensation, and proprioception, as well as 10 types of non-neuronal cells. Cross-species integration, spatial transcriptomics, and microneurography enabled cell-type-specific comparisons of soma size and conduction velocity between species. Human DRG somata are larger across all cell types than their mouse counterparts, and the conduction velocities of human hair follicle innervating A-fibers are faster than in mice, suggesting a functional shift in rapid mechanical detection in humans. This integrated human DRG reference cell atlas provides a resource for exploring new molecular and physiological features of human DRG, which could help identify new strategies for treating chronic pain and other diseases of the peripheral nervous system.

## INTRODUCTION

The fundamental capacity to perceive touch, temperature, body position, itch and pain depends on the diverse sensory neurons found in the peripheral somatosensory ganglia, including the 31 pairs of dorsal root ganglion (DRG) [1, 2] and one pair of trigeminal ganglion (TG) [3]. The importance of this system is underscored by the debilitating consequences of its dysfunction, including balance and coordination deficits and chronic pain [4]. Peripheral sensory neurons have been historically classified into broad categories based on conduction velocity, soma size, and a limited repertoire of molecular markers [5]. Large-diameter, fast-conducting neurons are termed A-fibers, and unmyelinated, slow-conducting neurons are termed C-fibers. A-fibers tend to be specialized for but not exclusively sensitive to mechanical stimuli, whereas C-fibers can detect a broader range of environmental stimuli including thermal, mechanical, and chemical stimuli.

Recent advances in single-cell transcriptomics have resolved at least 20 molecularly specialized neuronal subtypes in mice [2, 6–9], most of which have been shown to represent functionally distinct populations with unique nerve terminal morphologies, soma sizes, non-neuronal interactions, electrophysiological and stimulus response properties [2, 10]. This granular classification has provided key mechanistic insights into sensory encoding in both physiological and pathological states such as neuropathic pain [11–14]. Similar efforts have begun to molecularly and physiologically profile human DRG neurons [9, 15–20], but these efforts have been constrained by limited sample sizes, anatomical scope, and the inherent sparsity of neurons present in each ganglion. Thus, the DRG neuronal subtypes present in human and their relationship to mouse DRG neurons remains largely unclear.

Here, we overcome these limitations by constructing a reference atlas of the human DRG from 126 donors across the lifespan that spans cervical, thoracic, and lumbar levels. This atlas combines high-depth single-soma/cell RNA-sequencing (scRNA-seq) with large-scale single-nucleus RNA-sequencing (snRNA-seq) to resolve 22 neuronal subtypes, including several previously unrecognized molecular subtypes implicated in nociception, mechanosensation, thermosensation, and proprioception, and 10 non-neuronal subtypes. We validate the anatomical organization and prevalence of these subtypes using spatial transcriptomics and systematically identify their mouse counterparts. These data reveal that human DRG somata are substantially larger than their mouse counterparts across all neuronal subtypes. Recordings of human A-fiber low-threshold mechanoreceptors innervating hair follicles exhibit faster conduction velocities compared to their counterparts in mice. This study defines the molecular and cellular architecture of human DRG and supplies a foundational reference and tool (http://humanDRG.painseq.com) for developing targeted therapies for peripheral nervous system disorders.

## RESULTS

### The human DRG atlas identifies 22 neuronal subtypes

We constructed a comprehensive human DRG atlas by generating and integrating scRNA-seq and snRNA-seq (Fig. 1A) data from 126 human donors (Fig. 1B; Table S1). This cohort spans three spinal levels (cervical, thoracic, lumbar), both sexes, and ages from infancy to 90 years (Fig. 1C). We used both high-throughput snRNA-seq and deeply sequenced whole-soma RNA-sequencing [20] to construct the largest human DRG atlas to date (53,596 neurons and 498,444 non-neurons; Fig. 1B, Fig. S1A). By profiling nearly 4-fold more donors and 20-fold more neurons than currently available data [9], this new reference atlas provides a view of the human DRG cellular landscape.

**Figure 1.**
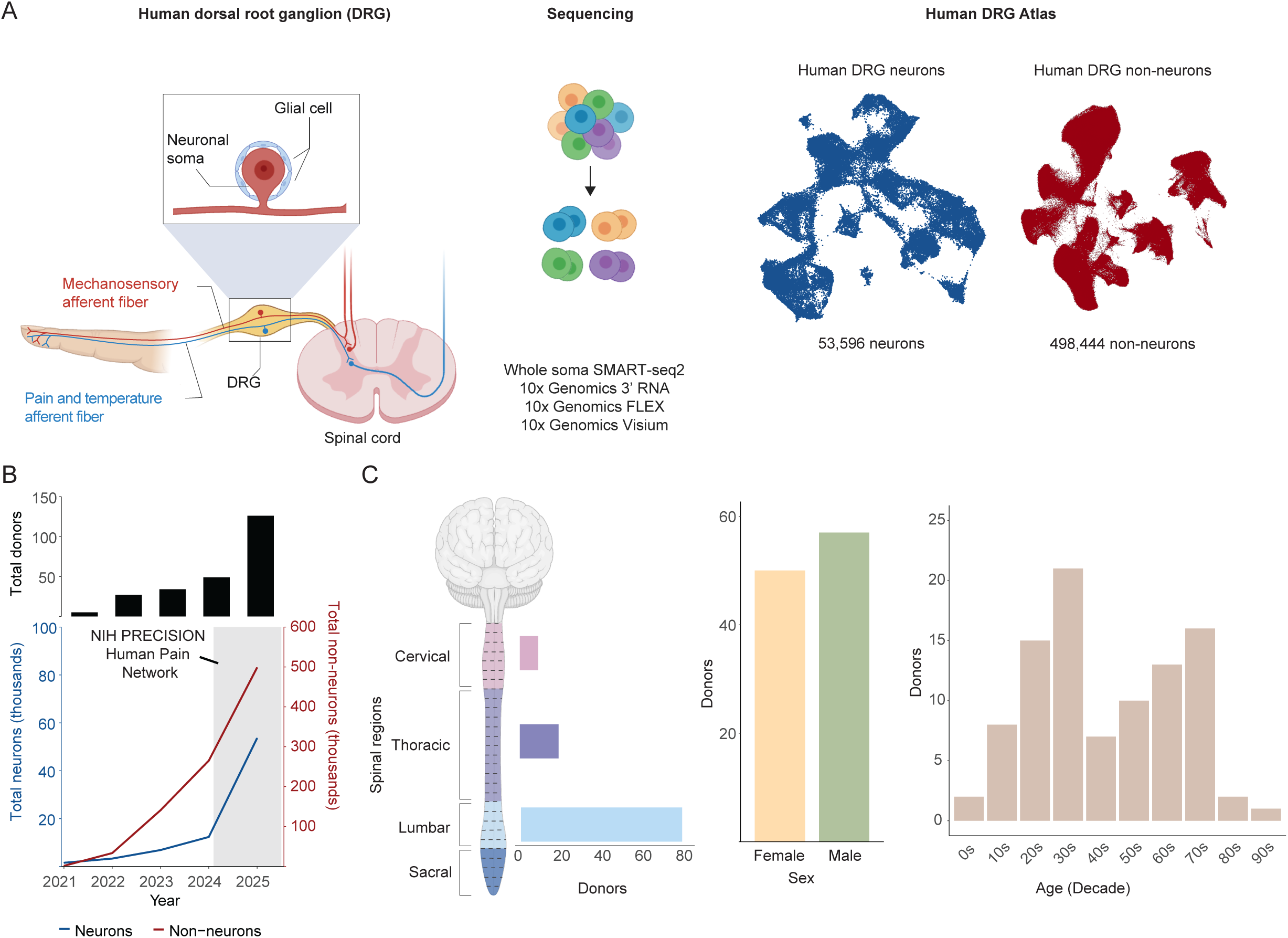
The human DRG cell atlas. (A) Left: Diagram of DRG anatomy and major cell types. Center: Dissociated human DRG cells and nuclei were profiled using SMART-seq2, 10x Genomics 3’ v3, and 10x Genomics FLEX platforms. Right: UMAP plots describing integrated DRG cell atlases comprising 53,596 neurons and 498,444 non-neuronal cells. (B) Top: bar plot displaying the number of DRG donors from which sc/snRNA-seq was performed since 2021, Bottom: line plot displaying the number DRG neurons (blue) and non-neurons (red) profiled by sc/snRNA-seq since 2021. Data from this study is shaded. (C) Bar plots displaying the number of donors included in the DRG cell atlas, grouped by the spinal level (Left), sex (Center), or age (Right). Donors without a recorded spinal level or age were excluded from this plot.

Within this integrated dataset we identified 22 transcriptionally distinct neuronal subtypes (Fig. 2 A-D). We assigned each subtype to A- or C-fiber using a small, predefined marker set for fast-conducting, myelinated neurons, specifically genes associated with the node-of-Ranvier and myelination (Fig. S1B–C). Twelve subtypes were annotated as A-fibers, nine as C-fibers, and one unclassified (ATF3, see methods). This A- and C-fiber classification is consistent with the hierarchical clustering of neuronal transcriptomes, which revealed that the primary transcriptional division among all neuronal subtypes corresponds to their A- or C-fiber designation (Fig. 2C). Additionally, an A-fiber-specific hdWGCNA module of 120 genes (Fig. S1D) [10] and *in situ* measurements (Fig. S1E-F) further corroborated this fiber-type split. We then annotated subtypes using a standardized scheme: primary class (A or C), functional family based on marker genes from previous studies, and one or two discriminative marker genes.

**Figure 2.**
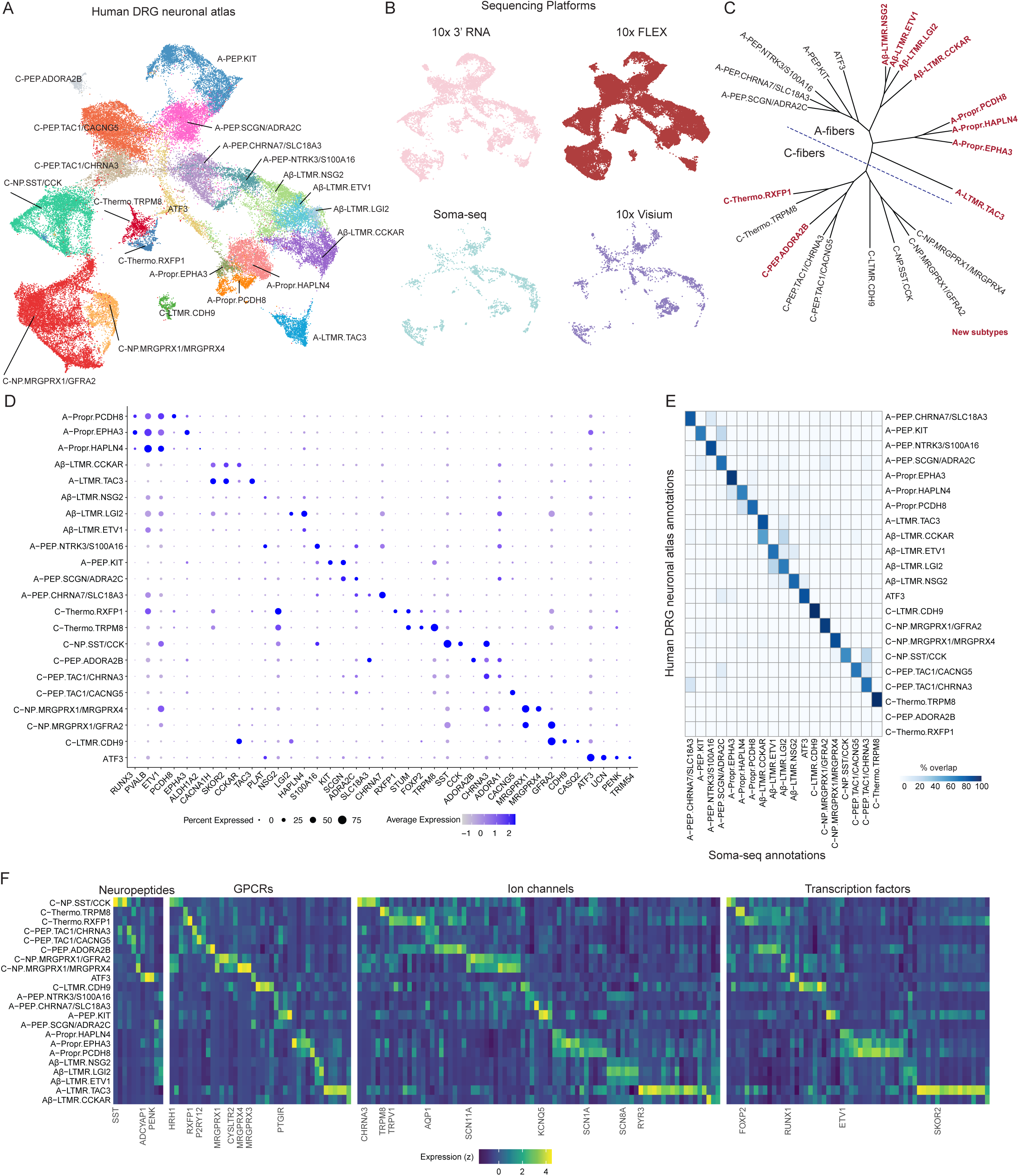
Transcriptional diversity of 22 human DRG neuronal subtypes. (A) Human DRG neuronal atlas (53,596 cells/nuclei) projected into UMAP space. Cells/nuclei are colored by their final cell type annotations. (B) UMAP projection displays the human DRG neuronal atlas, colored by sequencing platform (10x Genomics 3’ RNA: 9,125 nuclei; 10x Genomics FLEX: 40,917 nuclei; Soma-seq: 1,598 cells; and 10x Visium: 1,956 cells). (C) Dendrogram displays the hierarchical relationship of human sensory neurons. This hierarchal relationship was calculated based on the Euclidean distance of the principal components. Branch lengths are scaled to Euclidean distance. Dotted line indicates division between A-fibers and C-fibers. New cell types identified in this resource are colored in red. (D) Dot plot displays cell-type-specific marker gene expression in DRG neuronal subtypes. Dot size indicates the fraction of cells/nuclei expressing each gene, and color indicates average log-normalized scaled expression of each gene. (E) The heatmap displays the percentage of subtype annotations in the human DRG neuronal atlas that correspond to those independently identified in the soma-seq datasets (see Methods). Neuronal classifications differed for three subtypes: Aβ-LTMR.CCKAR was divided into two clusters in the human DRG neuronal atlas, while C-PEP.ADORA2B and C-Thermo.RXFP1 were identified only in the human DRG neuronal atlas. (F) The heatmap displays genes classified as neuropeptides, GPCRs, ion channels, and transcription factors that are enriched in each neuronal subtype relative to all other neuronal subtypes (log,FC > 0.75, adj. p<0.05). For visualization, only the top 15 genes (or less) are displayed per subtype per classification.

The 12 A-fiber subtypes include three *PVALB+* and *RUNX3+* proprioceptors (A-Propr.HAPLN4, A-Propr.EPHA3, A-Propr.PCDH8; Fig. 2D and Fig. S1G), four *NTRK3+* and *NTRK2+* Aβ-low-threshold mechanoreceptors (LTMRs) subtypes (Aβ-LTMR.ETV1, Aβ-LTMR.LGI, Aβ-LTMR.NSG2 and Aβ-LTMR.CCKAR), one A-LTMR (A-LTMR.TAC3), and four *CALCA*-, *TAC1-* and *NTRK1-*positive polymodal peptidergic subtypes (A-PEP.NTRK3/S100A16, A-PEP.KIT, A-PEP.SCGN/ADRA2C, and A-PEP.CHRNA7/SLC18A3). These classes align with prior human DRG reports [9, 20], however our new dataset resolves new subtypes within proprioceptors and Aβ-LTMRs (Fig. 2C).

We also identified nine C-fiber subtypes, including a C-LTMR subtype (C-LTMR.CDH9); three *CALCA*-, *TAC1-* and *NTRK1-*positive polymodal peptidergic nociceptors (C-PEP.TAC1/CHRNA3, C-PEP.TAC1/CACNG5, C-PEP.ADORA2B); two MRGPRX-expressing polymodal non-canonical peptidergic (NP) nociceptors (C-NP.MRGPRX1/GFRA2, C-NP.MRGPRX1/MRGPRX4); a distinct NP SST-positive subtype (C-NP.SST/CCK); and two temperature-sensitive populations (C-Thermo.TRPM8 and C-Thermo.RXFP1). These C-fiber subtypes align with prior human DRG reports [9, 20].

As most neuronal subtypes were detected by all sequencing platforms (Fig. 2E and Fig. S1H), we took advantage of the transcriptional depth of single soma-seq with the breadth of nuclei from snRNA-seq and next identified genes enriched in each neuronal subtype relative to all other neurons. We found that the 22 subtypes were enriched (log,fc > 0.75, adjusted (adj.) p-value < 0.05 relative to other neurons) for 1,896 genes (Table S2). These included therapeutically relevant gene families such as 11 neuropeptides (e.g., *ADCYAP1*), 40 GPCRs (e.g., *MRGPRX3*), 80 ion channels (e.g., *CACNA1E*), and 104 transcription factors (e.g., *SKOR2*; Fig. 2F). Collectively, these data describe cell-type-specific expression of lowly expressed genes, achieved through approximately 5 log,-fold greater transcriptomic depth than previous studies.

### The human DRG atlas identifies 10 non-neuronal populations and their interactions with neurons

Non-neuronal DRG cells play crucial roles in regulating the neuronal microenvironment, modulating neuronal physiology through direct interactions, and responding to insults [21, 22]. In the human DRG, we identified 10 major non-neuronal cell types across 498,444 cells/nuclei (Fig. 3A-B). These cell types include satellite glial cells (SGC), myelinating Schwann cells (mySC), non-myelinating Schwann cells (nmSC), endothelial cells, mural cells, epineurial cells (Fibroblast-Epi), perineurial cells (Fibroblast-Peri), endoneurial cells (Fibroblast-Endo), adipocytes, and immune cells. These cell types were annotated based on marker genes identified in previous human and mouse snRNA-seq studies [9, 21, 23–27].

**Figure 3.**
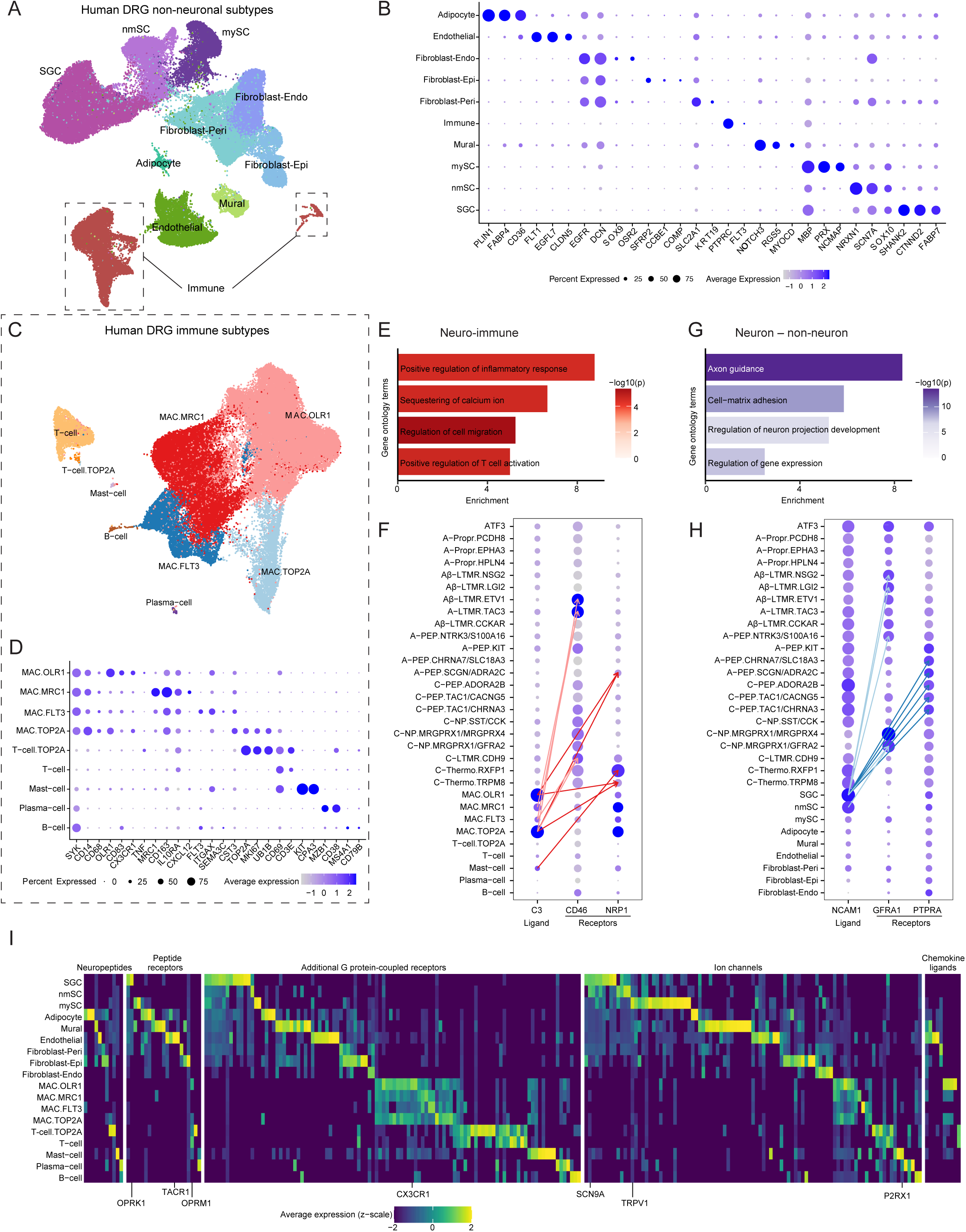
Non-neuronal cells in the human DRG. (A) Human DRG non-neuronal atlas projected into UMAP space (498,444 nuclei). Nuclei are colored by cell types. (B) Dot plot displays cell-type-specific marker gene expression in DRG non-neuronal subtypes. Dot size indicates the fraction of nuclei expressing each gene, and color indicates average log-normalized scaled expression of each gene. (C) Human DRG immune cells projected into UMAP space (59,469 nuclei). Nuclei are colored by immune subtypes. (D) Dot plot displays the expression of select marker genes for immune subtypes. Dot size indicates the fraction of nuclei expressing each gene, and color indicates average log-normalized scaled expression of each gene. (E) and (G) Bar plots display the GO terms enriched in genes that exhibit significant ligand-receptor interactions between neurons and immune cells (E) or neurons and other non-neuronal cells (G) compared to all other ligand-receptor pairs. (F) Dot plot displays the gene expression of ligand *C3* and its receptors *NRP1* and CD46 in individual neuronal and immune subtypes. Dot size indicates the fraction of nuclei expressing each gene, and color indicates average log-normalized scaled expression of each gene. The arrows link the top 5 (ranked by aggregated rank) cell type pairs for each ligand-receptor interaction. (G) Dot plot displays the gene expression of ligand *NCAM1* and its receptors, including *GFRA1* and *PTPRA*, in individual neuronal subtypes and non-neuronal cell types. Dot size indicates the fraction of nuclei expressing each gene, and color indicates average log-normalized scaled expression of each gene. The arrows link the top 5 (ranked by aggregated rank) cell type pairs for each ligand-receptor interaction. (H) Heatmap displays genes that are differentially expressed (log_2_FC>1, FDR<0.05) in each non-neuronal subtype relative to all other non-neuronal subtype. For each gene, cell types with expression in less than 5% of the nuclei are colored dark blue.

We subclustered the immune cells and identified nine subtypes, including B cells, T cells, mast cells, plasma cells, and macrophages based on established marker genes (Fig. 3C-D). We observed four transcriptionally distinct clusters of macrophages, largely consistent with prior reports [21]. Although all four macrophage clusters express pan-macrophage markers, such as *SYK*, *CD68* and *CD14*, each macrophage cluster exhibits distinct gene expression. *OLR1*+ macrophages express pro-inflammatory markers, such as *OLR1*, *CD83, CX3CR1, and TNF*, whereas *MRC1*+ macrophages express anti-inflammatory markers, such as *MRC1*, *CD163*, *IL10RA*, and *CXCL12* [28–31]. *FLT3+* macrophages express marker genes associated with tissue recognition and cell migration, such as *FLT3, ITGAX, SEMA3C, and CST3* [32, 33]. We also identified macrophage and T cell populations expressing proliferation markers, including *TOP2A, MKI67* and *BUB1B*, likely corresponding to the self-renewal populations previously reported [31].

Neuro-immune and neuro-glial interactions in the DRG are increasingly recognized as key drivers of pathological pain [28, 29, 34–36].To better understand how cellular interactions may shape human DRG function and pain signaling, we mapped ligand-receptor (LR) interactions between 22 neuronal subtypes and 18 non-neuronal cell types and immune subtypes in the DRG. This analysis revealed 153 significant neuro-immune interactions and 324 neuron-non-immune interactions (Table S3). Gene ontology analysis predicted that neuro–glia interactions regulate inflammation, cell migration, and T cell activation (Fig. 3E–F), while neuron–non-neuron cell interactions support axon guidance and neurite outgrowth (Fig. 3G). These pathways are increasingly implicated in chronic pain and response to nerve injury, and these atlases have implicated the specific neuronal and non-neuronal subtypes involved. For example, C-Thermo.RXFP1 and C-Thermo.TRPM8 neurons expressing Neuropilin-1 (*NRP1*), a known axon guidance receptor also linked to pain sensitivity [37–39], are predicted to interact with macrophages via C3–NPR1 signaling (Fig. 3F). Other subtype-specific signaling modules, such as predicted NCAM1-based interactions with multiple receptor partners, further illustrate how axonal growth, cell adhesion, and neuron–glia communication may be differentially regulated across neuronal subtypes (Fig. 3H).

Additionally, we examined physiologically relevant genes, including those encoding neuropeptides, receptors, and ion channels, and identified 243 genes differentially expressed across DRG non-neuronal cell types (log_2_fc>0.5, adjusted p-value<0.05 relative to all other non-neurons; Fig. 3I, Table S4). Together, these neuronal and non-neuronal atlases provide a comprehensive cellular and molecular map of the human DRG, resolving 22 neuronal and 10 non-neuronal subtypes. The atlas reveals previously unrecognized A-fiber cell types, rare C-fiber populations, and novel neuro-immune and neuro-glial signaling pathways, establishing a foundation for studying human somatosensory biology and pain mechanisms.

### Spatial transcriptomics reveals molecular and structural subtype diversity of human DRG neurons

To investigate the spatial distribution and soma sizes of transcriptionally defined neuronal subtypes in human DRG, we profiled lumbar DRGs using two high-resolution spatial transcriptomics platforms: MERSCOPE and Xenium (Fig. S2A). The MERSCOPE panel included 300 genes, and the Xenium panel included 472 genes, with 50 genes overlapping between the two panels (Table S5). All neuronal subtypes from the human DRG neuronal atlas were detected in the spatial transcriptomics data (Fig. 4A-B; S2B-C), and neuronal subtype-specific marker gene expression (log_2_fc>0.5, adj. p-value < 0.05 compared to other neurons; Fig. 4C) was positively correlated (Spearman’s r > 0.8) between spatial data and the human DRG neuronal atlas, with the exception of A-LTMR.TAC3.

**Figure 4.**
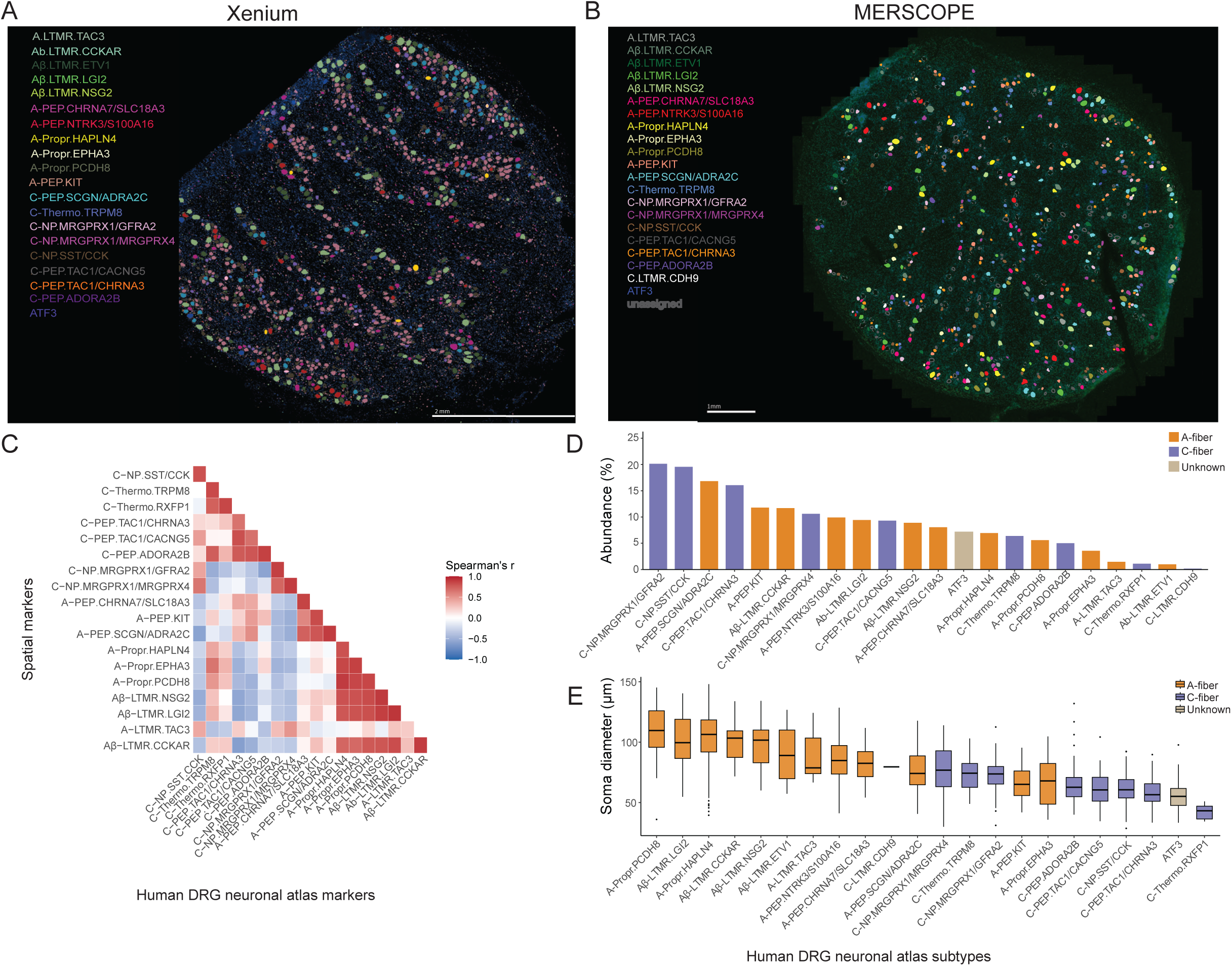
Spatially defined DRG neuronal subtypes. (A) and (B) Representative Xenium (A) or MERSCOPE (B) spatial transcriptomics images of human DRG. Segmented neuronal regions are colored by their transcriptionally defined neuronal subtype. Scale bar: 2 mm (Xenium) and 1 mm (MERSCOPE). (B) Heatmap displays correlations between subtype specific markers from the DRG neuronal atlas and the spatial transcriptomic datasets (merged Xenium and MERSCOPE). Heatmap is colored by Spearman’s *r*. (C) Bar plot displays percentage of neuronal subtypes detected in each spatial platform. Xenium and MERSCOPE data were merged. Bars are colored by fiber class. Xenium (1186 cells from 1 donor) and MERSCOPE (619 cells from 1 donor) data were merged. (D) Box plot displays soma diameters of each neuronal subtype. Boxes are colored by fiber class. Xenium (1186 cells from 1 donor) and MERSCOPE (619 cells from 1 donor) data were merged.

Knowledge of cell type abundances provides practical insight into the feasibility of studying specific neuronal populations, as rare subtypes can present technical and analytical challenges. Consistent with previous human sequencing studies, the most abundant subtypes are C-NP.MRGPRX1/GFRA2, C-NP.SST/CCK, and A-PEP.SCGN/ADRA2C, while the rarest subtypes are C-Thermo.RXFP1 Aβ-LTMR.ETV1, and C-LTMR.CDH9 (Fig. 4D). The abundance of neuronal subtypes largely correlates (Spearman’s *r* = 0.76) between the human DRG neuronal atlas and the spatial transcriptomics data. This strong concordance between the integrated and spatial datasets underscores the reliability of the human DRG neuronal atlas for quantifying and comparing neuronal subtype abundances within the ganglion.

Soma size is an important correlate of the degree of myelination, conduction velocity and function, so we next measured the diameter for each of the 22 neuronal subtypes in the spatial transcriptomics data (Fig. 4E). Soma sizes of A-fibers and C-fibers were consistent with the myelination scores (Fig. S1C) and hierarchical classification (Fig. 2C). Similar to previous spatial transcriptomic surveys of the human DRG [18, 20], we observed larger diameters (median range: 90-100 µm) in Aβ-LTMR and proprioceptor subtypes, intermediate sizes (median range: 81-89 µm) in A-PEP neurons, and smaller diameters (median range: 42-80 µm) in C-fiber and ATF3-expressing subtypes.

Similar to mouse DRG neurons, human A-fiber somata are generally larger than those of C-fibers, with mouse Aδ-LTMRs measuring 25–40 µm, Aβ-LTMRs 40–54 µm, and proprioceptors >55 µm, whereas C-fiber neurons are typically <25 µm. However, we identified two notable exceptions in humans: A-PEP.KIT (median: 69 µm) and A-Propr.EPHA3 (median: 67 µm) exhibited smaller soma sizes compared to other human proprioceptors and Aβ-LTMRs, falling within the size range observed with human C-fibers. Our measurements for A-Propr.EPHA3 align with previous observations in mice, where specific proprioceptor subtypes (II_1,2,4_) that innervate muscle spindle afferents also exhibit smaller diameters compared to Type Ia and Type Ib proprioceptors and other A-fibers [40]. Finally, a mouse A-PEP population with a diameter comparable to mouse C-fibers has not been identified.

Overall, these spatial transcriptomics data provide orthogonal evidence for each of the transcriptionally defined subtypes identified in the human DRG neuronal atlas and add important information about neuronal morphology within the human DRG.

### Neuronal subtype differences between DRG spinal levels

As different body regions have specialized somatosensory requirements, we next examined how human DRG neuronal subtype composition varies across spinal levels. To determine how the proportion of neuronal subtypes vary across spinal levels, we fit a beta regression model with spinal level as the primary predictor, adjusting for age, sex, institution, tissue collection method and sequencing platform (Fig. S3A; Table S6), and applied *post hoc* pairwise comparisons to the fitted model. These comparisons revealed that 7 subtypes had significantly different proportions across spinal regions (Fig. 5A). Mechanosensory A-fiber populations, including two proprioceptor subtypes (A-Propr.HAPLN4 and A-Propr.EPHA3; Fig. 5B), were significantly enriched in cervical DRGs compared to thoracic and lumbar regions (Fig. 5B; Fig. S3B). In contrast, the proportions of several nociceptor and thermosensory subtypes, including C-NP.MRGPRX1/MRGPRX4, C-Thermo.RXFP1, C-PEP.TAC1/CHRNA3, A-PEP.SCGN/ADRA2C, and C-NP.SST/CCK, were higher in thoracic or lumbar DRGs compared to cervical DRGs. These trends reflect known functional specialization along the spinal axis, from proprioceptive control of the upper body [40] to visceral and lower-limb sensory input [2, 41, 42].

**Figure 5.**
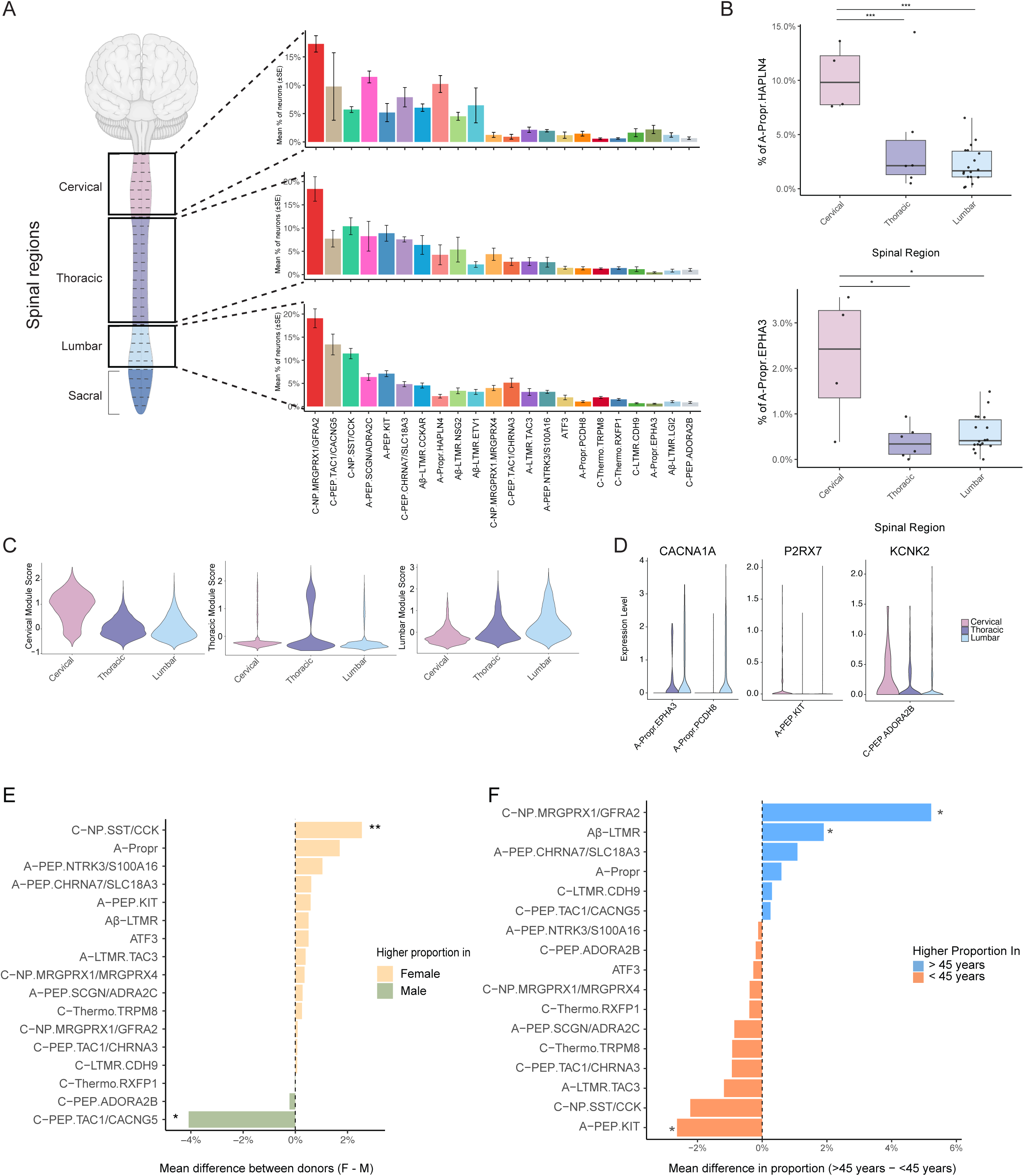
Variation in neuronal subtype composition across spinal levels, sex, and age. (A) Bar plots display proportional distributions of neuronal subtypes across in, thoracic, or lumbar DRGs (± SEM across donors). Bars are colored by neuronal subtype and ordered by median abundance across all spinal regions. (B) Box plots display the proportions of A-Propr.HAPLN4 and A-Propr.EPHA3 subtypes in cervical, thoracic, and lumbar DRGs. Each dot represents one donor. Donors with <1,000 neuronal nuclei profiled by sc/snRNA-seq were excluded. Significance was assessed using a beta regression model (*: p < 0.05; **: p < 0.01; ***: p < 0.005). (C) Violin plots of gene module scores associated with DRG neurons from each spinal level. Modules were defined by genes enriched in all neurons from a given region relative to other regions (log,FC > 1; adj. p < 0.05; Table S7). (D) Violin plots display log-normalized expression levels of *CACNA1A*, *P2RX7*, and *KCNK2*, separated by spinal region (cervical, thoracic, or lumbar). (E) Bar plot displays the mean difference (female – male) in subtype proportions between sexes. Donors with <1,000 neuronal nuclei were excluded. Bars are colored by sexes. F: female; M: male; *: p < 0.05; **: p < 0.01. (F) Bar plot displays the mean difference in neuronal subtype proportions between donors younger than 45 years and those 45 years or older. Donors with <1,000 neuronal nuclei profiled by sc/snRNA-seq were excluded. Bars are colored by age (<45 or >45). *: p < 0.05.

Level-specific transcriptional differences between neuronal subtypes may also contribute to functional specialization, so we compared neuronal gene expression patterns between cervical, thoracic, and lumbar DRGs. We first compared neurons at each spinal level using a pseudobulk differential expression approach where we controlled for sex, age, sequencing technology, and number of cells/nuclei (see methods). For each spinal region, we identified a core module of 9 to 13 genes consistently enriched (Fig. 5C; log_2_FC>1; adj. p-value<0.05; Table S7) compared to all other regions, including canonical developmental regulators such as *HOXA1*. These findings are consistent with the well-established role of *HOX* genes in specifying the distinct segmental identities of neurons along the spinal cord [43, 44]. In addition to these shared patterns, individual subtypes exhibited distinct level-specific gene expression signatures (Table S8). For instance, *CACNA1A* was more highly expressed in lumbar and lower thoracic DRG for A-Propr.EPHA3 and A-Propr.PCDH8 subtypes when compared to their counterparts at cervical levels (Fig. 5D; log_2_FC>1; adj. p-value<0.05). *CACNA1A* encodes for the alpha subunit of the calcium channel CAV2.1, a known regulator of action potential of muscle spindle afferents [45]. Together, these findings suggest that both the proportions and gene expression profiles of human DRG neuronal subtypes are regionally tuned to support the specialized sensory functions of different spinal levels.

### Modest sex- and age-associated shifts in DRG neuronal composition

As differences in pain and sensory perception have been reported between sexes [18, 46–48] and across lifespan [49–51], we next asked whether the proportion of DRG neuronal subtypes or their gene expression patterns are affected by sex or age. For each subtype, we fit two separate beta regression models: one testing for sex differences and one for age group differences (<45 vs. ≥45 years), each adjusting for spinal region, institution, sequencing platform, surgical vs. post-mortem DRG collection and either age or sex (Table S6). To account for A-fiber subtypes that were either rare or inconsistently represented within each age group, functionally related A-fiber classes (A-Propr and Aβ-LTMRs) were combined prior to modeling to improve statistical robustness. Post hoc comparisons were used on the fitted model to identify significant differences between groups (Table S6).

The sex-stratified model revealed that most neuronal subtypes are observed in similar proportions between males and female and across age (Figs. S3B-C), with only C-NP.SST/CCK and C-PEP.TAC1/CANCG5 displaying subtle but statistically significant sex differences in their proportions (Fig. 5E), and C-NP.MRGPRX1/MRGPRX4, Aβ-LTMRs, and A-PEP.KIT displaying subtle but statistically significant age differences in their proportions (Fig. 5F). Notably, despite the well-documented decline in tactile sensitivity with age [52], we observed a slight increase in the proportion of Aβ-LTMRs in DRGs from donors > 45 years old than < 45 years old. Age-related changes in cutaneous end organ innervation rather than Aβ-LTMR loss per se may contribute to these age-related changes in tactile sensitivity [53]. Overall, these findings suggest that sex and age have a measurable but limited impact on neuronal subtype composition in the human DRG.

We next asked whether there are gene expression differences in DRG neuronal subtypes between sexes or across age. Between sexes, we identified 211 genes that were upregulated or downregulated in females compared to males (log_2_FC>1; adj. p-value<0.05; Table S9), including *NAV3,* a gene responsible for neuronal morphogenesis and whose intronic SNPs have been associated with post-operative chronic pain [54]. Across ages, we identified 1,451 genes upregulated or downregulated in older individuals compared to younger individuals (Table S10), including the ryanodine receptor-cation channel *RYR2*, which has been implicated in nociceptor hyperexcitability and neuropathic pain states in obese mice [55].

### Transcriptional similarity of human and mouse sensory subtypes

Comparing transcriptomes between human and mouse DRG neuronal subtypes remains critical for translating mechanistic insights from animal models to human biology. To compare subtype-specific transcriptomes across species, we examined transcriptomic similarities between human and mouse subtypes by (1) integrating our new human DRG neuronal atlas with a recently published scRNA-seq atlas of seven mouse datasets [10] (Fig. S4A-B), (2) trained a variational autoencoder (VAE) on the recently published scRNA-seq mouse atlas to predict cell type identities of human clusters (Fig. S4C-E) and (3) trained a label transfer algorithm to a new harmonized atlas of 21 mouse sc/snRNA-seq studies to classify each human cell or nuclei as a mouse cell type (Fig. S4F-H).

Across all analytical approaches, we identified largely consistent transcriptomic relationships between human and mouse DRG subtypes. Hierarchical clustering of the integrated human–mouse dataset, based on Euclidean distances between cluster centroids across the top 30 principal components, grouped human C-LTMR.CDH9 most closely with mouse C-LTMRs (Fig. 6A). Both the VAE model (Fig. 6B) and the label transfer analysis (Fig. S4F–H) supported this relationship. Similar to C-LTMR.CDH9, most human subtypes showed consistent cross-species relationships across all methods, however the A-LTMR.TAC3 subtype produced varying results. Hierarchical clustering positioned A-LTMR.TAC3 apart from all human and mouse A-fiber populations, the VAE model linked it to the mouse peptidergic C-fiber population PEP1.1, and label transfer classified most A-LTMR.TAC3 cells/nuclei as mouse Aδ-LTMRs. This pattern may reflect a human-specific mechanoreceptor subtype or indicate that additional A-fiber-enriched sequencing in mouse is needed to resolve this population.

**Figure 6.**
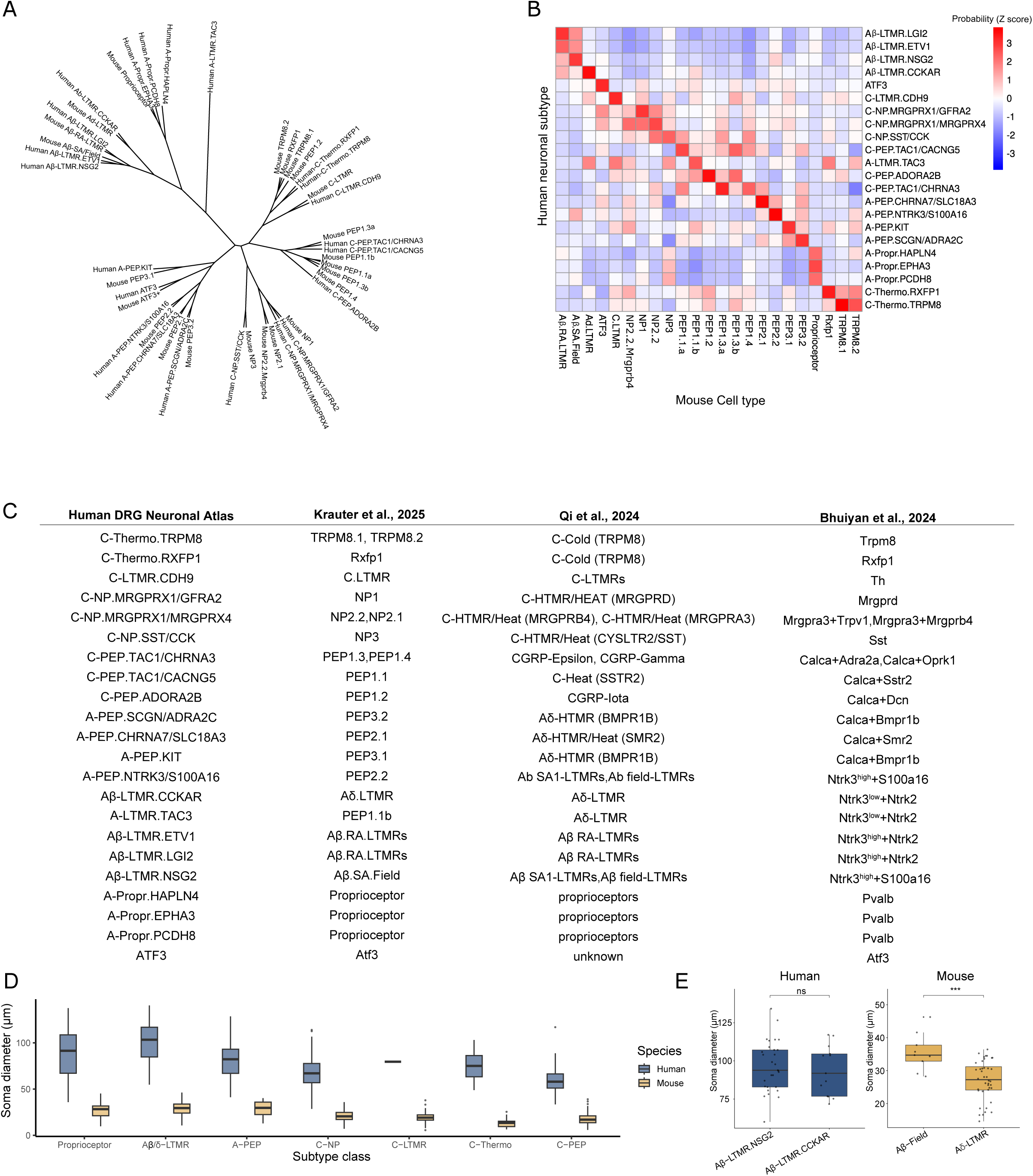
Cross-species analysis of mouse and human DRG neurons. (A) Dendrogram displays cell type similarities of mouse and human DRG neurons after cross-species integration by canonical correlation analysis. This hierarchical relationship was calculated based on the Euclidean distance of the principal components. Branch lengths are scaled to Euclidean distance. (B) Heatmap displays z-scored cell type classification probabilities from the variational autoencoder model between mouse (columns) and human (rows) cell types. (C) Table displays human DRG neuronal subtypes with their corresponding mouse DRG neuronal subtypes. For each human cell type (rows), we report the corresponding mouse cell type based on the nomenclature reported by Krauter et al. (2025), Qi et al. (2024), and Bhuiyan et al. (2024). (D) Box plot displays the distribution of somata sizes per neuronal subtype class for human and mouse. Human somata sizes are calculated from spatial transcriptomics data presented in Figure 4, while mouse data are calculated from the recent Krauter et al., 2025 study. Boxes are colored and split by species, and subtypes are merged into physiological classes. (E) Box plot displays distribution of soma sizes for Aβ-LTMR.NSG2 and Aβ-LTMR.CCKAR, along with their transcriptomic correlate in mouse, Aβ-Field and Aδ-LTMR. Human somata sizes are calculated from spatial transcriptomics data presented in Figure 4, while mouse data are calculated from the recent Krauter et al., 2025 study. Boxes are colored by species. Statistical significance was calculated using a Wilcoxon rank sum test (p-value > 0.05 = “n.s”, p-value < 0.001 = “***”).

Based on the results from the human-mouse integration, the VAE classifier and the label transfer, we outline the DRG neuron subtype correlates between human and mouse, including different nomenclature schemes (Fig. 6C; Table S12).

### Species differences in conduction velocity among human hair follicle LTMRs with direction selectivity

Using the DRG spatial transcriptomics data above (see Fig. 4), we next compared the soma sizes of human DRG neuronal classes to their counterparts in mouse. In addition to the observation that soma sizes of all human DRG neuronal classes are larger than their counterparts in mouse (Fig. 6D), we noted that the soma sizes of A-LTMR subtypes are more similar in human than they are in mice. Specifically, in mice, the median soma sizes of A-LTMR subtypes range from 34.7 µm to 27.3 µm, with Aδ-LTMR soma sizes significantly smaller than Aβ-LTMRs (Wilcoxon test *P <* 0.001; Fig 6E). In contrast, the soma sizes of human A-LTMRs range from 97.1 µm to 91.9 µm, with the Aδ-LTMR transcriptomic equivalent (Aβ-LTMR.CCKAR), exhibiting comparable soma sizes to other human Aβ-LTMR subtypes (Wilcoxon test *P* > 0.5). To explore whether these species differences in A-LTMR soma size correspond to differences in their electrophysiological properties, we performed microneurography recordings from individual somatosensory afferent fibers (units).

One hundred and eighty-three LTMRs (upper limb, n=135; lower limb, n=48), all sensitive to gentle touch (soft brushing), were recorded in awake human participants. A and C fibers were distinguished based on spike morphology and response latency. A-LTMRs were further classified by their adapting properties during ramp-and-hold indentation into rapidly and slowly adapting types, following established criteria (Figure S5) [56, 57]. Among rapidly adapting A fibers, hair follicle afferents (A-LTMR(HF)s) responded to single-hair deflection and light air puffs, while Field-LTMRs displayed multiple spots of high sensitivity within the receptive field and did not respond to either hair displacement or remote tapping of the skin. Slowly adapting units comprised SA1 and SA2 types, distinguished primarily by their discharge pattern during sustained indentation – SA1s showing irregular firing and SA2s regular firing. Among these A-LTMRs, conduction velocities were similar across subtypes (Fig. 7A). A-LTMR(HF)s responded to hair movement and air-puff stimulation, and the air-puff response was abolished after shaving the receptive field (Fig. 7B–C). Further, A-LTMR(HF)s showed directional tuning to brushing in proximal-distal versus distal-proximal directions, with directional preferences ranging up to several-fold (Fig. 7D– E). Direction preference was defined by a ≥20% higher mean firing frequency in one direction compared to the opposite, consistent across brush velocities and stable within each unit. In contrast to A-LTMR(HF)s, C-LTMRs – the only other type of human DRG afferent known to respond to hair movement (Fig. 7F) [58] – displayed no directional preference to brushing (Fig. 7G-H).

**Figure 7.**
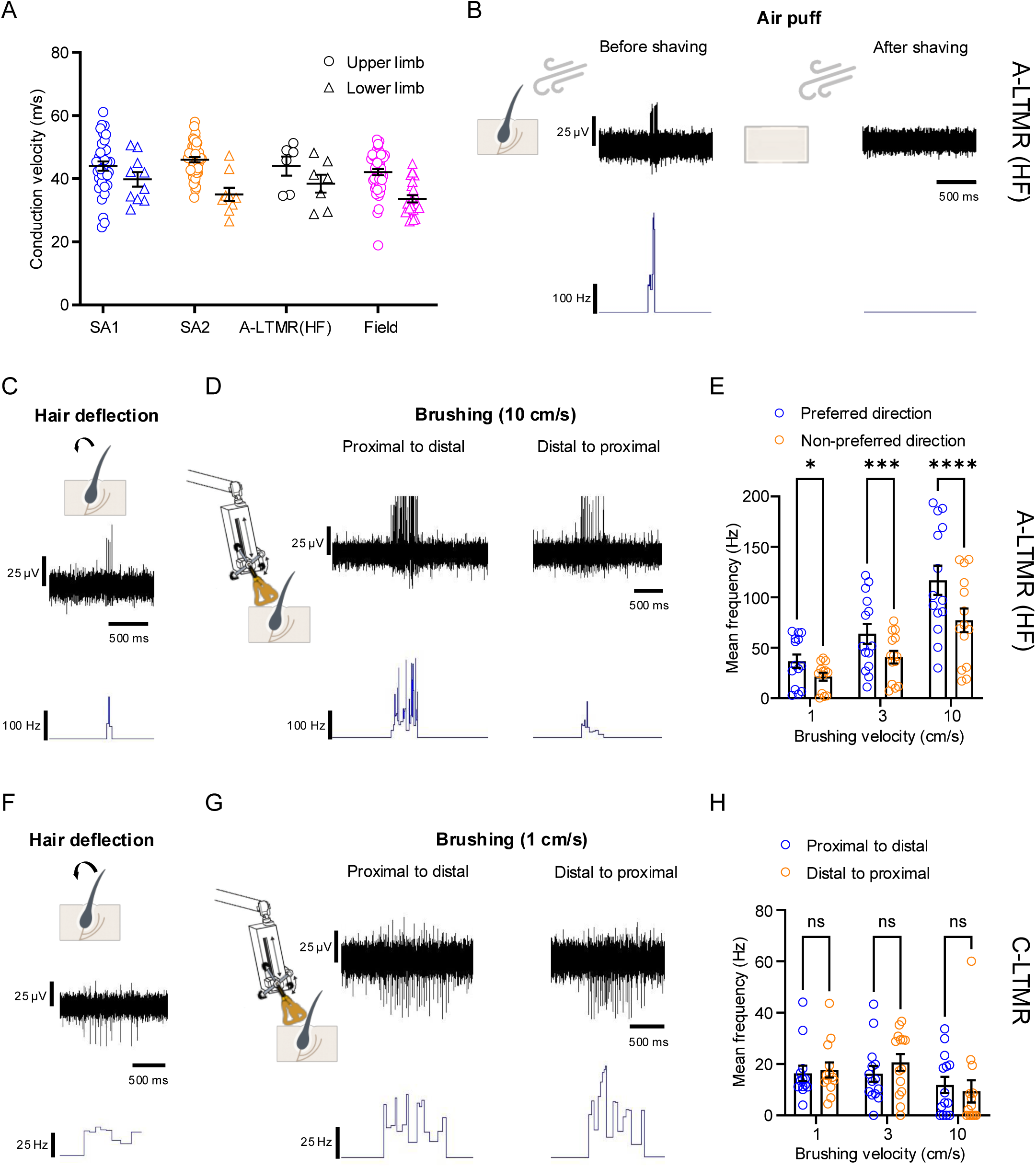
Physiological properties of human A-LTMR subtypes. (A) Plot displays the recorded conduction velocities of SA1, SA2, A-LTMR(HF), and Field receptors (mean ± SEM). Each point represents an individual neuron; circles, upper limb (n=130 neurons); triangles, lower limb (n=48 neurons). Conduction velocities did not differ among subtypes in the upper limb (Kruskal–Wallis, P = 0.0922) or lower limb (P = 0.0836). HF: hair follicle. (B) Top left: Traces display representative A-LTMR(HF) spike activity in response to an air puff. Bottom left: Trace displays instantaneous spike frequency. Top right: Trace displays spike activity recorded after shaving the receptive field. Bottom right: Trace displays instantaneous spike frequency (C) Traces display representative A-LTMR(HF) spike activity in response to single-hair deflection (CV = 40.2 m/s). Top: Traces display raw spike trace; Bottom: Trace displays instantaneous spike frequency. (D) Top: Traces display representative A-LTMR(HF) spike activity in response to brushing at 10 cm/s in both directions. Bottom: Trace displays instantaneous spike frequency. (E) Box plot displays mean firing frequencies of A-LTMR(HF)s to brushing at 1, 3, and 10 cm/s, grouped by directional preference. For each recorded unit, brushing directions (proximal-distal and distal-proximal) were reassigned as preferred or non-preferred, with preference defined as a ≥20% higher mean firing frequency than the opposite direction across velocities. Direction preference was stable within each unit. Each circle represents an individual trial (n=3 neurons, multiple trials per neuron). Responses were significantly greater in the preferred direction (mixed-effects model, P < 0.0001). Mean ± SEM., n.s.: not significant. *P < 0.05, ***P < 0.001, ****P < 0.0001. (F) Traces display representative C-LTMR spike activity in response to single-hair deflection. Top: Traces display raw spike trace; Bottom: Trace displays instantaneous spike frequency. (G) Top: Traces display representative C-LTMR spike activity in response to brushing at 1 cm/s in both directions. Bottom: Traces display application of stimuli, in both directions. (H) Bar plot displays mean firing frequencies of C-LTMR to brushing at 1, 3, and 10 cm/s, comparing proximal-distal and distal-proximal strokes. Anatomical directions are shown, as no consistent direction preference was observed across units. Each circle represents an individual trial (n=5 neurons, multiple trials per neuron). C-LTMR responses were not modulated by brushing direction (2-way RM-ANOVA, P = 0.6501). Data are mean ± s.e.m.; n.s.: not significant.

Taken together, these physiology recordings, including evidence of directional tuning, indicate that human A-LTMR(HF)s are functionally analogous to mouse Aδ-LTMRs. In mice, Aδ-LTMRs specifically innervate awl hairs and exhibit direction-selective responses to caudal-to-rostral deflection. This property is not observed in mouse Aβ or C-LTMRs [59]. Notably, however, human A-LTMR(HF) conduction velocities lie in the Aβ (Ab) range, similar to SA1, SA2, and field LTMRs, potentially indicating conserved function but an upward shift in conduction velocity of this type of A-LTMRs from mice to humans. Cross-species transcriptomic analysis suggests that the mouse Aδ-LTMR is transcriptionally most similar to the human Aβ-LTMR.CCKAR.

The connection between SA- and Field-LTMR physiology and specific transcriptomic subtypes remains unresolved. Cross-species transcriptomic analysis identifies Aβ-LTMR.NSG2 as the likely correlate Aβ-SA/Field-LTMRs in mice, but future work is needed to clarify how the physiological diversity of Aβ-SA and Field LTMRs aligns with their comparatively similar transcriptomes in both species.

## DISCUSSION

We constructed a reference atlas of the human dorsal root ganglion (DRG) by integrating snRNA-seq, soma-seq, and spatial transcriptomics from 126 donors across cervical, thoracic, and lumbar levels. This publicly available resource (http://humanDRG.painseq.com) describes both known sensory neuron classes and resolves previously unrecognized human subtypes spanning nociception, mechanosensation, thermosensation, and proprioception. Additionally, spatial transcriptomics confirmed each of these subtypes *in situ* and clarified their soma sizes, which are larger across all neuronal subtypes than their mouse counterparts. In addition to their larger soma sizes, there is also an acceleration in conduction velocity for at least the hair follicle innervating A-LTMRs compared to those in mice. Together, this single-cell and spatial transcriptomic resource provides a detailed characterization of human DRG cell types and enables cross-species comparisons that can advance both mechanistic and translational research on peripheral sensory system disorders.

### Functional predictions of human somatosensory neuron subtypes

Cross-species transcriptomic comparisons between human and genetically defined mouse subtypes provide an opportunity to generate hypotheses about the corresponding human DRG neuronal subtype function.

#### C-Thermoreceptors (C-Thermo)

Temperature-sensitive neurons help organisms detect both innocuous and noxious temperature shifts in the environment [60]. Mouse and human datasets both contain two closely related C-Thermo subtypes: a population expressing the cold-sensing ion channel *TRPM8* [61, 62] and a population expressing the relaxin receptor *RXFP1* [60, 63], indicating broad conservation of temperature-sensitive neuron classes across species. Both the human C-Thermo.RXFP1 and the mouse Rxfp1 subtype share high expression of *TRPV1* and *TRPM2* [9, 10], consistent with a response profile in the warmth range.

Notable species differences emerge within the TRPM8-lineage neurons. In humans, the C-Thermo.TRPM8 cluster shows substantial co-expression of *TRPV1* with *TRPM8* (∼70%), suggesting unresolved heterogeneity that may include both unimodal cool-sensing and bimodal cold/heat-responsive neurons. In contrast, mouse scRNA-seq datasets show greater molecular diversification, with distinct Trpm8.1 (cool-only) and Trpm8.2 (cool- and heat-responsive) subtypes in addition to *Rxfp1*. Physiological studies of human C-fibers responsive to both menthol/cold and heat support the existence of this mixed functionality in the human population [20, 64].

#### C-Peptidergic neurons (C-PEP)

Classically known as peptidergic nociceptors, C-PEP neurons were first characterized as *CGRP, Substance P*, and *NtrkA* positive polymodal neurons that respond to high heat and mechanical force. Human and mouse C-PEP neurons show strong transcriptional correspondence, reflecting a conserved class of peptidergic nociceptors involved in detecting noxious stimuli. The human C-PEP.TAC1/CACNG5 population aligns with the mouse PEP1.1 heat-sensitive nociceptors (C-heat neurons) [2]. C-PEP.ADORA2B shows a one-to-one correspondence with PEP1.2 (Calca+Dcn), maintaining expression of *TRPV1*, *NTRK2*, and *ADORA2B* but lacking *PIEZO2*, consistent with a dedicated C-heat-responsive role [10]. C-PEP.TAC1/CHRNA3 subtype exhibits molecular features of both mouse PEP1.3 and PEP1.4, which in mice are linked to deep-tissue and visceral innervation as well as high-threshold mechanosensitivity and noxious heat [2, 42]. Since PEP1.3 is more abundant in mouse DRGs with visceral innervation, sampling additional human spinal levels may better resolve transcriptomic substructure within C-PEP.TAC1/CHRNA3.

#### C-Non-canonical Peptideric neurons (C-NP)

Classically referred to as non-peptidergic nociceptors, C-NP neurons constitute a polymodal class that detects heat and high-threshold mechanical stimuli (C-MH) as well as diverse noxious and pruritogenic cues. As some human NP neurons express similar neuropeptides as C-PEP, we repurposed the acronym to improve accuracy while facilitating comparison to prior studies. Cross-species mapping supports one-to-one alignment of human subtypes with mouse groups: C-NP.MRGPRX1/GFRA2 = NP1, C-NP.MRGPRX1/MRGPRX4 = NP2, and C-NP.SST/CCK = NP3, indicating a conserved organizational logic for these classes of polymodal C-fibers. We also observe sex-associated differences in C-NP.SST/CCK and age-related shifts in C-NP.MRGPRX1/MRGPRX4, which could relate to variation in pain thresholds and itch sensitivity [65], although targeted studies are needed.

We observed a notable species difference in the NP2 class. Mouse NP2 divides into NP2.1(Mrgpra3+Trpv1) and NP2.2 (Mrgpra3+Mrgprb4), but we did not resolve an analogous subdivision within human C-NP.MRGPRX1/MRGPRX4 in this study, despite subclustering the C-NP.MRGPRX1/MRGPRX4.

#### C-low-threshold mechanoreceptors (C-LTMR)

C-LTMRs are specialized sensory neurons in hairy skin that detect slow, gentle stroking, which is crucial for the perception of pleasant touch. In both species, a conserved C-LTMR population is present that expresses CDH9 as a marker gene. While mouse studies have described C-LTMRs as enriched in upper compared to lower spinal levels, we did not observe a significant difference across spinal levels. This may reflect the fact that the human DRG neuronal atlas includes only the cervical level C2 and does not include thoracic DRGs above T10 (see limitations).

#### ATF3

In both mouse and humans, ATF3 neurons are a rare and perplexing class of neurons that may correspond to a functionally distinct cell type or a simply a transient cell state of other neuronal subtypes. As a group, these cells have small soma sizes (Fig. 4E) despite being transcriptionally more similar to A-fibers than C-fibers. ATF3 neurons express genes that are associated with nerve regeneration (e.g. *ATF3, SPRR1A*) [12], suggesting that ATF3 neurons could represent multiple neuronal subtypes in a shared regenerative state. However, rare populations of ATF3 neurons have been observed in apparently uninjured animals across multiple species and sensory ganglia, which could also be consistent with a rare but stable cell type.

#### A-Peptidergic neurons (A-PEP)

A-PEPs are fast-conducting nerve fibers that initiate the reflexive or protective responses to potentially damaging stimuli [20]. A-PEPs are high-threshold polymodal Aβ or Aδ-fibers and can terminate in the skin as either free or circumferential endings in the skin [66]. We identified four human A-PEP subtypes with one-to-one correspondence to mouse A-PEP subtypes: A-PEP.CHRNA7/SLC18A3 corresponds to PEP2.1 (Calca+Smr2); A-PEP.NTRK3/S100A16 corresponds to PEP2.2 (Ntrk3^high^+S100A16); A-PEP.KIT corresponds to PEP3.1 (Calca+Bmpr1b); and A-PEP.SCGN/ADRA2C corresponds to PEP3.2 (Calca+Bmpr1b).

#### A-Proprioceptors (A-Propr)

Responsible for our sense of body position, proprioceptors are classically divided into three functional types (Ia, Ib, and II) [40]. Type Ia innervates muscle spindles and detects stretch velocity, Type Ib innervates the Golgi tendon organ and detects muscle tension, and Type II innervates muscle spindles and detects static muscle position. While mouse genetic tools have described molecular profiles of each proprioceptor subtype, we have comparatively limited insight into these subtypes in human [40, 67, 68]. Our dataset now overcomes this challenge by resolving three distinct human proprioceptor types that correspond to Ia, Ib, and II proprioceptor subtypes. Cross-species analysis indicates that A-Propr.HAPLN4 likely corresponds to Type Ia proprioceptors, A-Propr.PCDH8 likely corresponds to Type Ib proprioceptors, and A-Propr.EPHA3 correspond to Type II proprioceptors.

A-Propr.HAPLN4 and A-Propr.EPHA3 neurons are more abundant in cervical DRGs compared to thoracic and lumbar levels. In mouse, proprioceptor subtypes also show level-specific enrichment, but this typically occurs within the cervico-brachial region (C5–C8 in mouse; C5–T1 in human) [40]. In contrast, all cervical DRGs analyzed in the present study are derived from C2, which innervates suboccipital and deep dorsal neck muscles. These muscles also contain a high density of muscle spindles, essential for monitoring head position, gaze stabilization, and posture [69].

#### A-low-threshold mechanoreceptors (A-LTMRs)

Aβ- and Aδ-low-threshold mechanoreceptors (Aβ-LTMRs/Aδ-LTMRs) mediate discriminative touch, classically divided into seven physiological subtypes by their nerve end morphology, response properties and conduction velocity [70]. Single-cell transcriptomic studies have identified comparatively fewer A-LTMR subtypes in both mouse (three subtypes) and human (five subtypes), suggesting that some physiological differences between LTMRs may not be fully explained by their differences in gene expression alone.

Three of the transcriptionally defined human A-LTMR subtypes have 1:1 transcriptional correlates with mice: (1) Aβ-LTMR.LGI2 corresponds to Aβ-rapidly adapting LTMRs (Aβ-RA-LTMRs), which respond to dynamic indentation and vibration in hairy and non-hairy skin; (2) Aβ-LTMR.NSG2 corresponds to Aβ-slowly adapting/field-LTMRs (Aβ-SA/Field-LTMRs), which respond to skin stretch and wide-field brushing, respectively; and (3) Aβ-LTMR.CCKAR corresponds to Aδ-LTMRs, which detect gentle, directionally tuned hair deflection. The other two human LTMR transcriptomic subtypes are more difficult to align with mouse LTMRs. Human Aβ-LTMR.ETV1 shows similarity to both mouse Aβ-Field/SA-LTMRs and Aβ-RA-LTMRs [71], whereas human A-LTMR.TAC3 corresponds to different sets of A-LTMR subtypes depending on the computational approach used.

In addition to transcriptomic differences between human and mouse LTMRs, we also observed that the conduction velocity of human direction-tuned hair-follicle A-LTMR units conduct predominantly within the Aβ range, whereas in mice they conduct within the Aδ range. Together, LTMRs in humans and mice exhibit a broadly conserved organizational logic for encoding tactile stimuli, yet key transcriptomic and physiological divergences highlight evolutionary refinements that may underlie species-specific differences in mechanosensation.

### Evolutionary conservation and divergence of DRG subtype function

While human and mouse DRG cell types are broadly conserved at the transcriptomic level, we observed specific gene expression (Table S11), morphological and physiological differences. Human neurons have larger somata across classes (Fig. 5D), and because soma size correlates closely with axon caliber and degree of myelination, larger cells are expected to conduct more rapidly. Soma size also increases with species body size [22] and our measurements are consistent with this scaling. In line with these principles, LTMR subtypes exhibited higher conduction velocities in humans than in mouse. These findings are consistent with human HTMR conduction velocities [20, 58], and suggest a general pattern of faster peripheral signaling in humans than mice. The reasons for this interspecies difference are unclear, but accelerated conduction velocities in humans may preserve temporal precision over longer distances than is required in rodents.

The physiological and anatomical differences observed here illustrate how evolutionary pressures may fine-tune molecular features of broadly conserved sensory neuron subtypes to meet species-specific demands. Comparative analyses across the nervous system reveal that, although many neuronal subtypes are conserved, individual genes often show species-specific expression patterns that may underlie functional divergence between corresponding subtypes [3, 9, 19, 72]. With a comparable number of cells/nuclei, subtypes, and sequencing depth to the mouse atlases [2, 9, 10], the human DRG atlas presented here provides a foundation for identifying subtype-specific gene expression differences between species.

### Limitations of the Study

Although our beta regression model helped adjust for high donor to donor variability, the model’s overall explanatory power was modest (median pseudo *R*^2^ of 0.44 across all subtypes). This indicates that a substantial portion of the per donor variability in neuronal composition remains unexplained and could be influenced by genetic background, environmental factors, or other unmeasured variables. Consequently, the observed associations between cell type proportions and factors such as spinal level, sex, or age should be confirmed in in larger and more diverse cohorts.

Correlations between molecularly defined cell types and neuronal physiology and function largely rely on cross-species homology with genetically defined studies in the mouse. While this is a powerful approach, innervation targets may differ across species or within subtypes of the same species, which may limit this approach. It is also likely that some physiological differences cannot be captured by transcriptomic methods alone such that additional measurement methods (e.g. channel phosphorylation states) are needed to fully achieve 1 to 1 correspondence between physiologically-defined and molecularly-defined neuronal subtypes.

In the spatial transcriptomics datasets, some cell types annotations were based on the absence of marker genes. While our spatial annotations largely aligned with more rigorous annotations by sc/snRNA-seq (Fig. 4C), limitations in probe detection rates may introduce some error in spatial annotations.

Finally, our anatomical sampling was limited to specific cervical, thoracic, and lumbar spinal levels, which could overlook cell type specialization in the unsampled DRGs.

### Conclusions

We constructed a reference atlas of the human dorsal root ganglion by integrating single-nucleus, soma, and spatial transcriptomic data from 126 donors. The atlas resolves 22 neuronal and 10 non-neuronal subtypes, each mapped back to the ganglion in situ. We systematically compare these subtypes across spinal levels, sexes, age groups, and species. Cross-species analyses reveal both deep conservation and meaningful divergence in DRG organization, including larger somata and faster conduction velocities in human neurons compared to mouse. These features point to evolutionary adaptations in the structure and physiology of sensory neurons that may enhance conduction speed and temporal precision in humans. Beyond cataloging cell types, this resource provides a foundation for linking molecular identity to function, refining experimental models, and developing strategies to selectively target human sensory neuron subtypes in the treatment of chronic pain and other peripheral nervous system disorders.

## METHODS

### Ethical compliance and declaration

#### University of Pennsylvania

This study complies with all relevant ethical regulations and the revised Declaration of Helsinki. The protocol for collecting DRG tissues from consented patients was approved by the University of Pennsylvania IRB (IRB834222). The National Disease Research Interchange (NDRI, supported by the National Institutes of Health (NIH) grant U42OD11158) has an IRB protocol (IRB704541) for procuring human tissues from postmortem donors, approved by the University of Pennsylvania IRB committee. The Luo laboratory research application was approved by the NDRI Feasibility Committee (RLUW1 01). In addition, as determined by the University of Pennsylvania IRB committee, the Luo laboratory experiments using human DRG samples from de-identified consented postmortem donors were exempted from the human subject requirements.

#### Linköping University

The protocol of performing microneurographic recordings in consenting human participants was approved by the Swedish Ethical Review Authority (Dnr 2015/305-31, 2017/485–31, and 2020-04426) and complied with the revised Declaration of Helsinki. Exclusion criteria included neurological or musculoskeletal disorders, skin diseases (e.g., psoriasis), diabetes, and recent use of pain-relieving or psychoactive medication.

#### Harvard Medical School

All procedures for human tissue procurement and ethical oversight were reviewed and approved by the Institutional Review Board at Brigham and Women’s Hospital [2024P002201/MGB1924]. Protocol MGB1924 was reviewed by Mass General Brigham IRB and determined to be not human research. DRGs were collected from consented donors using a rapid autopsy protocol at Mass General Brigham. All donors were deidentified.

#### University of Texas at Dallas

All procedures for human tissue procurement and ethical oversight were reviewed and approved by the Institutional Review Board at the University of Texas at Dallas (protocol Legacy-MR-15-237). Donors were also prospectively recruited at the University of Washington (IRB #10916) and MD Anderson Cancer Center (2013-0871).

##### Organ Donors

DRGs were collected from organ donors through a partnership with the Southwest Transplant Alliance, an organ procurement organization (OPO) based in Texas. The Southwest Transplant Alliance, Anabios, and their OPO partners obtain informed consent for research tissue donation either directly from the donor (through first-person consent, such as a driver’s license or other legally binding document) or from the donor’s legal next of kin.

Donor screening and consent follow the guidelines set by the United Network for Organ Sharing (UNOS). In addition, OPOs comply with standards established by the US Centers for Disease Control and are inspected every two years by the Department of Health and Human Services. Medical information provided with donor tissue is shared in accordance with HIPAA regulations to ensure donor privacy is protected (Shier, S.I. 2025).

##### Surgical Donors

Patients aged 18 years or older undergoing C1–C2 arthrodesis for neck pain at the University of Washington were prospectively recruited (IRB #10916). Informed consent was obtained from all participants, who received a $125 gift card for participation. Patients were identified through surgeon notifications of scheduled procedures and daily electronic surgery schedules via the UW Medicine Enterprise Data Warehouse. Exclusion criteria included cognitive or language limitations that prevented informed consent. Eligible patients were approached either by phone (outpatients) or in person (inpatients), and consent was obtained using a standardized script. Pain was classified as acute (<3 months) or chronic (≥3 months) according to International Association for the Study of Pain (IASP) definitions [73].

Patients undergoing thoracic vertebrectomy for malignant spinal tumors at MD Anderson Cancer Center were recruited with informed consent. Demographic, clinical, and medical history data were obtained through retrospective review of medical records and prospective collection at the time of enrollment. Neuropathic or radicular pain was assessed for each dermatome according to the guidelines of the IASP Neuropathic Pain Special Interest Group (definite or probable neuropathic pain). During surgery, spinal nerve roots were ligated for reconstruction or tumor resection, and DRGs were collected as previously described [74].

#### Washington University in St. Louis

Human DRGs were obtained from organ donors with full legal consent for use of tissue in research and in compliance with procedures approved by Mid-America Transplant. The Human Research Protection Office at Washington University in St. Louis provided an Institutional Review Board waiver.

### Human tissue collection and sequencing

We generated human DRGs transcriptomic data using multiple complementary approaches across four centers. Tissues were obtained from postmortem organ donors and surgical donors with appropriate consent, and processed using laser capture microdissection (LCM), single-nucleus RNA-sequencing (snRNA-seq), or snMultiome profiling. Below, we detail the protocols used at each site for tissue handling, library preparation, sequencing, and data processing.

#### University of Pennsylvania (UPenn)

Human DRG tissues used in this study were procured through the NDRI. Tissue recovery was authorized and consented by the donor next of kin for research use (unpaid). DRGs were dissected from postmortem donors and immediately frozen in optimal cutting temperature (OCT) compound, and the frozen blocks were shipped in dry ice to the Luo laboratory. Upon arrival, the tissue blocks were stored at –80 °C until use. Information regarding DRGs, experiments performed, de-identified donors and screening criteria are summarized in Table S1.

##### LCM of human DRGs

The DRGs embedded in OCT were cryosectioned (Leica, CM1950 Cryostat) into 20-µm sections and mounted onto Arcturus PEN Membrane Frame Slides (Applied Biosystems, LCM0521). One of every five consecutive sections was collected for LCM to avoid repeated dissection of the soma from the same neuron in d ifferent sections. The slides were stored at –80 °C until further use.

On the day of LCM, the slides were transferred to the Skin Biology and Disease Resource-based Center LCM Core on dry ice. Before dissection, the section was briefly stained with Rnase-free Arcturus HistoGene staining solution (Applied Biosystems, 12241-05) for better visualization of neuronal soma: 70% cold ethyl alcohol (EtOH) for 30 s; HistoGene staining for 10 s; 70% cold EtOH for 30 s; 95% cold EtOH for 30 s; 100% cold EtOH for 30 s; and air dry for 2 min. Then, the slide was put onto an LCM microscope system (Leica, LMD6) for the neuronal soma dissection. The laser was calibrated, and the laser intensity was adjusted to achieve the best dissection efficiency. The dissected individual neuronal somata were collected in the cap of a 200-µl polymerase chain reaction (PCR) tube containing 4 µl of lysis buffer [20]. The sequencing library was generated following the Smart-seq2 workflow [20]. There were three quality control steps before the sequencing. (1) Housekeeping gene GAPDH and neuronal marker PGP9.5 were amplified after reverse transcription and pre-amplification. Samples showing clear, specific GAPDH and PGP9.5 bands were retained. (2) The concentration after the first purification was measured, and samples with concentrations greater than 0.2 ng/µl were selected. (3) The concentration of the final library was measured. Only libraries that passed all three quality control steps were submitted for sequencing.

The libraries were pooled together (384 samples for one batch) and sequenced on a NovaSeq 6000 platform at the Children’s Hospital of Philadelphia Center for Applied Genomics. Raw sequencing data were demultiplexed with bcl2fastq2 version 2.20 (Illumina) followed by Tn5 transposon adapter sequence trimming with Cutadapt (version 3.5). The processed reads were then aligned to the human genome (GRCh38 GENCODE as the reference genome and GRCh38.104. GTF as the annotation) using STAR version 2.7.9a49. STAR quantMode GeneCounts was used to quantify unique mapped reads to gene counts.

#### Single-nucleus RNA-sequencing (UTD)

Lumbar and thoracic DRGs were obtained from organ donors, while cervical and thoracic DRGs were collected from surgical donors for single-nucleus RNA-sequencing. Samples were rapidly frozen in pulverized dry ice immediately after recovery and stored at –80 °C until processing.

For nuclei isolation, frozen DRG tissue was finely minced (∼1 mm pieces) in chilled isolation buffer (0.25 M sucrose, 150 mM potassium chloride (KCl), 5 mM magnesium chloride (MgCl,), 1 M Tris pH 8.0, 0.1 mM DTT, protease inhibitor, 0.1% Triton X-100, and 0.2 U/µl Rnase inhibitor), followed by gentle homogenization on ice using a Dounce homogenizer (∼15 strokes). The homogenate was filtered through a 100 µm strainer and centrifuged (500 ×g, 10 minutes, 4 °C). The resulting nuclei pellet was resuspended in 2 ml of buffer (1% bovine serum albumin [BSA] in phosphate buffered saline [PBS], 0.2 U/µl Rnase inhibitor) and centrifuged again (500 ×g, 5 minutes, 4 °C). The pellet was immediately fixed using the 10x Genomics Chromium Fixed RNA Profiling kit (16 hours, 4 °C), followed by labeling with the 10x Fixed RNA Feature Barcode kit (16 hours). Subsequent library preparation followed the manufacturer’s instructions. A detailed version of the nuclei isolation and processing protocol is available at <u>protocols.io</u> [75]

Sequencing of libraries was performed at different batches and were sequenced on the Illumina NextSeq2000 at the University of Texas at Dallas Genome Core or NovaSeqX at Psomogen. The reads were processed and aligned to the human reference genome (GRCh38) using Cell Ranger v7 (10x Genomics). We ran the remove-background function of CellBender (v0.3.0) to remove ambient RNA and empty droplets from raw expression matrices using default parameters.

Prior to integrating data from all centers, we first separately annotated neurons and non-neurons using Seurat v4. Nuclei with more than 1,000 detected genes and less than 10% mitochondrial counts were retained. The ambient RNA corrected counts matrix for each library was normalized, 2,000 variable genes were identified, integration anchors were selected across all libraries, and the data were scaled. The number of principal components (PCs) was determined as the point where additional PCs contributed less than 5% of the standard deviation of highly variable gene expression, and the selected PCs cumulatively explained at least 90%. Libraries were integrated using Seurat’s canonical correlation analysis (CCA), and nuclei were clustered and embedded in UMAP space at a resolution of 0.5. Clusters enriched for *SNAP25* (log_2_FC > 0.5, adj. p-value < 0.05) were annotated as neurons, and those enriched for *SPARC*, *MPZ,* or *PTPRC* under the same criteria were annotated as non-neurons.

#### Single-nucleus RNA-sequencing (HMS)

Single nuclei of human DRGs were collected using a previously described gradient protocol at HMS (86). Briefly, human DRGs were initially pulverized on dry ice and ∼0.5 to 1 cm^3^ of powder was placed into homogenization buffer [0.25 M sucrose, 25 mM KCl, 5 ml of MgCl2, 20 mM tricine-potassium hydroxide (KOH; pH 7.8), 1 mM dithiothreitol (DTT), actinomycin (5 µg/ml), 0.04% BSA, and ribonuclease (Rnase) inhibitor (0.1 U/µl)] for ∼15 seconds on ice. After the brief incubation for 15 seconds on ice, samples were transferred to a Dounce homogenizer for an additional 10 strokes with a tight pestle in a total volume of 5 mL of homogenization buffer. IGEPAL was added to a final concentration of 0.32%, and five additional strokes were performed with the tight pestle. The tissue homogenate was then passed through a 40-µm filter and diluted 1:1 with a working solution. Nuclei extracted were layered onto an iodixanol gradient after homogenization and ultracentrifuged as previously described (86). After ultracentrifugation, nuclei were collected between the 30 and 40% iodixanol layers and diluted for microfluidic encapsulation of individual nuclei in barcoded droplets.

Nuclei were labeled with DAPI to inspect their integrity under a fluorescent microscope and determine their quantity manually using a hemocytometer. Approximately, 10,000 nuclei were used for each ATAC and gene expression library. Libraries were generated according to the Chromium Next GEM single-cell multiome ATAC + gene expression guidelines (10x Genomics). Both ATAC and RNA libraries were shipped to Genewiz (Azenta Life Sciences) and were sequenced based on the sequencing parameters suggested by 10x genomics on the Illumina NovaSeq6000 platform.

Nuclei suspensions were sequenced using 10x Genomics assays, resuspended, and loaded into 10x Chromium device for snRNA-seq (10x Genomics v3.1). Libraries were prepared for snRNA-seq according to the manufacturer’s protocol. Libraries were sequenced on an Illumina NextSeq 500. Sequencing data were processed and mapped to the human (GRCh38) genome using 10x Genomics cellranger v7. We ran the remove-background function of CellBender (v0.3.0) to remove ambient RNA and empty droplets from raw expression matrices using default parameters.

Prior to integrating data from all centers, we first separately annotated neurons and non-neurons using Seurat v4. Nuclei with more than 1,000 detected genes and less than 10% mitochondrial counts were retained. The ambient RNA corrected counts matrix for each library was normalized, 2,000 variable genes were identified, integration anchors were selected across all libraries, and the data were scaled. The number of PCs was determined as the point where additional PCs contributed less than 5% of the standard deviation of highly variable gene expression, and the selected PCs cumulatively explained at least 90%. Libraries were integrated using Seurat’s CCA, and nuclei were clustered and embedded in UMAP space at a resolution of 0.5. Clusters enriched for *SNAP25* (log_2_FC > 0.5, adj. p-value < 0.05) were annotated as neurons, and those enriched for *SPARC*, *MPZ,* or *PTPRC* under the same criteria were annotated as non-neurons.

To maximize cost and efficiency, snRNA-seq was performed on batches of 1-13 donors. We used Souporcell to demultiplex sequencing reads to the level of individual donors [76]. Briefly, Souporcell detects genetic variants within sequenced RNA, generates a cell by variant genotype matrix, and clusters cells based on their allelic assignments. Souporcell version 2.5 was run using default settings. Clusters were initialized with known genotypes (imputed from DNA sequencing) provided by the –known_genotypes option. As input, we combined single cell RNA runs containing the same donors. We verified that samples contained the expected donors by cross-correlating cluster genotypes for each sample against imputed genotypes for all donors. We restricted analysis to Souporcell clusters whose genotypes were correlated with imputed genotypes with a Pearson correlation coefficient of greater than 0.7.

For genotyping, DNA purification was performed with up to 25mg of tissue using the QIAamp Fast DNA Tissue Kit and sent to Azenta Life Sciences for 4X WGS. Imputation was performed using GLIMPSE2 [77] and the HGDP-1kGP reference panel [78]. Only common variants (minor allele frequency > 5%) were used for Souporcell cluster initialization.

#### Washington University in St. Louis (WashU)

Human DRGs were obtained from organ donors with full legal consent for use of tissue in research and in compliance with procedures approved by Mid-America Transplant. The Human Research Protection Office at Washington University in St. Louis provided an Institutional Review Board waiver. Extraction and collection of human DRG tissues were performed as previously described in collaboration with Mid-America Transplant [48] with the following modifications and specifications. DRGs (T11 – L5) were surgically removed from postmortem organ donors, within 1 – 3 hours of aortic cross-clamping. Extracted tissues were immediately placed in ice-cold, oxygenated N-methyl-D-glucamine (NMDG)-based artificial cerebrospinal fluid (aCSF; 93 mM NMDG, 2.5 mM KCl, 1.25 mM NaH2PO4, 30 mM NaHCO3, 20 mM HEPES, 25 mM glucose, 5 mM ascorbic acid, 2 mM thiourea, 3 mM Na+ pyruvate, 10 mM MgSO4, 0.5 mM CaCl2, 12 mM N-acetylcysteine; adjusted to pH 7.3 using NMDG or HCl, and 300 – 310 mOsm using H2O or sucrose) and transported to the lab. Tissues were inspected, and adjacent tissues were removed. Cleaned tissues were snap-frozen in a liquid nitrogen vapor-based CryoPod Carrier before being transferred to -80 °C for long-term storage. To prepare for sequencing, human DRG tissues were embedded in OCT compound and sectioned on a cryostat. Sections were stored at -80 °C if not used immediately.

snMultiome-seq data of human DRGs were retrieved from a previous manuscript [79]. In summary, to test for the neuronal yield, single-nuclei resuspension was prepared using several protocols: a previously described gradient centrifugation protocol [80], a previously described non-gradient crude extraction protocol [24], and a commercial nuclear extraction kit (10x Genomics). Some samples also underwent fluorescence-activated cell/nucleus sorting (FACS) to remove cellular debris. We found that the gradient method yielded the highest neuronal percentage compared to other protocols, and that FACS negatively affected the neuronal coverage. Following the nuclear extraction, nuclei were diluted to target 10k nuclei recovery for each library. Diluted nuclei were further processed and prepared for sequencing according to the manufacturer’s manuals of 10x Genomics Chromium Single Cell Multiome Assay. Libraries were sequenced on an Illumina NovaSeqX Plus targeting 150,000 paired reads/nucleus for snRNA-seq libraries. Raw sequencing data from individual libraries were processed using 10x Genomics cellranger-arc (v2.0.1) and mapped to human reference genome GRCh38. We ran the remove-background function of CellBender (v0.3.0) on the cellranger generated counts matrices to remove ambient RNA and empty droplets from raw expression matrices using default parameters.

Prior to integrating data from all centers, we first separately annotated neurons and non-neurons using Seurat v4. Nuclei with more than 1,000 detected genes and less than 10% mitochondrial counts were retained. The ambient RNA corrected counts matrix for each library was normalized, 2,000 variable genes were identified, integration anchors were selected across all libraries, and the data were scaled. The number of PCs was determined as the point where additional PCs contributed less than 5% of the standard deviation of highly variable gene expression, and the selected PCs cumulatively explained at least 90%. Libraries were integrated using Seurat’s CCA, and nuclei were clustered and embedded in UMAP space at a resolution of 0.5. Clusters enriched for *SNAP25* (log_2_FC > 0.5, adjusted P < 0.05) were annotated as neurons, and those enriched for *SPARC*, *MPZ,* or *PTPRC* under the same criteria were annotated as non-neurons.

### Previously published human DRG studies

Prior studies have profiled human DRGs at cell-type resolution [9, 17–19, 24]. In a previous harmonized atlas study, we aggregated publicly available human sc/snRNA-seq datasets and generated additional human snRNA-seq data [9]. Raw reads were mapped to the human reference genome, and ambient RNA correction was performed. The resulting corrected count matrices served as input for constructing the integrated human reference atlas described in this study.

### Nomenclature

We developed a scalable nomenclature based on the multiple lines of evidence (Fig. S5). The name for each neuronal subtype is composed of three parts, systematically layering classical physiology with modern transcriptomics. The first component, fiber type, denotes the fundamental division between fast-conducting, myelinated ‘A’-fibers and slow-conducting, unmyelinated ‘C’-fibers. The second component, the “functional class”, describes the neuron’s primary cutaneous sensory function, which we inferred from known human or homologous mouse physiology. We use established abbreviations such as “Propr” for proprioceptors, “LTMR” for low-threshold mechanoreceptors (touch), “Thermo” for unimodal thermoreceptors (temperature), and PEP/NP for peptidergic/non-canonical peptidergic high-threshold mechanoreceptors (often polymodal/pain). This distinction for NP neurons reflects a key divergence from the mouse, as the human DRG neuronal atlas confirms that virtually all human C nociceptors express canonical peptidergic markers like CGRP and TrkA. For A-LTMRs, if the conduction velocity can be determined or estimated, a ‘β’ or ‘δ’ should follow the ‘A’. The final component is one to two molecular features, key gene markers (e.g., *KIT*, *TRPM8*, *PCDH8*) that uniquely define the cluster. This provides a stable transcriptomic anchor for the subtype, ensuring clarity for future studies even as we identify more granular subtypes. Thus, a name like A-Propr.HAPLN4 systematically describes a myelinated (A), proprioceptive (Propr) neuron that is uniquely identified by its expression of HAPLN4 compared to other A-Propr subtypes. This nomenclature is designed to honor the rich history of sensory neurobiology while providing a precise and scalable framework for future discoveries.

In mice, C-fiber nociceptors are divided into peptidergic and non-peptidergic subclasses based on the expression CGRP, TRPV1, Substance P and TrkA. In humans, all C fiber nociceptors express CGRP, TRPV1, and TrkA, and most express Substance P. However, we retain the NP nomenclature for cross-species homology but refer to these cells as **N**on-canonical **P**eptidergic to reflect their marker profile.

One key exception to the nomenclature is the cell type “ATF3”. This cell type has been identified in the naïve state in at least six species, however expresses the molecular features shared across DRG neurons after peripheral nerve injury, regardless of fiber type. As we cannot ascertain whether ATF3 cells are neurons in an injury state or a naïve cell type, we chose not to assign ATF3 with a fiber type.

### Construction of the human reference atlas

We integrated all neuronal data across centers and previously published datasets. The combined object was split by dataset or center, normalized, and variable genes were selected using the vst method, excluding mitochondrial genes (prefix “MT-”). Integration features and anchors were computed across subsets, and the data were integrated using Seurat’s CCA into an “integrated” assay, retaining the original “RNA” assay for marker visualization. The integrated object was scaled, PCA was performed, and PCs were selected using the same 90% cumulative and 5% marginal variance criteria. UMAP and t-SNE embeddings were computed, a shared nearest-neighbor graph was built, and clustering was performed at resolution 3. Violin plots (nFeature_RNA, nCount_RNA, percent.mt) were generated by Study for QC, and marker expression was visualized on UMAP and t-SNE for broad cell type markers (*SNAP25, RBFOX3, SPARC, MPZ, RBFOX1, PTPRC*) and curated neuronal subtype panels (Table S13). Doublets were removed if clusters showed enrichment for both neuronal and non-neuronal markers.

The non-neuronal atlas was constructed using the same workflow (res = 3), using non-neuronal markers as shown in Fig. 3B. Immune subclustering was performed on clusters enriched for PTPRC (CD45).

### Analysis of single-soma RNA-seq data of DRGs using Seurat and R

The single-soma RNA-sequencing dataset, which includes both the published dataset [20] and new generated dataset from the same three donors, comprises a total of 1,901 cells. For Seurat integration, the data were grouped into six virtual batches based on donor identity and sequencing run.

Initial Seurat v5 analysis revealed distinct clusters but two of them showed expression of non-neuronal markers (not shown). We interpreted these clusters as neurons contaminated with adjacent non-neuronal cells and excluded them from further analysis.

Clustering of remaining 1,785 cells reproduced all clusters in previous publication [20] and identified several new clusters. Firstly, proprioceptors were resolved into three distinct subclusters. Secondly, A-LTMRs formed seven clusters. However, three of them showed significantly fewer detected genes compared to other A-LTMR clusters and lacked specific marker genes. Similar features were also observed in hPEP.0 cluster described in previous publication [20]. With assumption that these clusters represent LCM technical artifacts (possibly sampling not central but side region of the cells), we excluded them (the three A-LTMR clusters and hPEP.0 cluster) from further analysis based on low gene detection relative to other clusters and absence of clear marker gene expression.

Final clustering in Seurat, performed on the remaining 1,391 cells, revealed a well-defined cluster structure. This structure closely recapitulated the previously published single-soma dataset [20], while new clusters emerged. The proprioceptor group now consisted of three distinct clusters. The A-LTMR group was divided by four distinct clusters.

### Cross-species analysis of human and mouse DRG neuronal subtypes

We first converted mouse genes in the mouse DRG atlas [10] to human orthologs using Ensembl’s biomart v102 corresponding to the GRCm38.p6 (mm10) reference genome assembly. We then selected genes with 1:1 matching homology between the species and a “Gene Order Conservation” score of >90. Next, we used Seurat v5 CCA to integrate the mouse data with the human data from this study. Within species, we merged the smallest data subsets to achieve a minimum of 30 cells per the lowest level of sample identity. We then processed the merged mouse-human dataset through the CCA integration by following the integration tutorial on the Seurat library website (https://satijalab.org/seurat/articles/seurat5_integration). We built the cluster dendrogram for the cross-species atlas using Seurat::BuildClusterTree (dims 1:30, reduction = “integrated.cca”) and calculated the pairwise distance matrix between cell type pairs from the dendrogram (using ape::cophenetic.phylo) for a heatmap visualization.

We used pollock [81] to build a variational autoencoder-based classifier using the humanized mouse DRG atlas [10]. For the training and testing with the mouse and human data (respectively) the Seurat objects were first transformed to python anndata using SeuratDisk (https://github.com/mojaveazure/seurat-disk). We built the classifier by following the package authors’ example notebook (https://github.com/ding-lab/pollock/blob/master/examples/pbmc_model_training_and_prediction.ipynb) using the default settings which produced a model with good quality metrics. For visualization as a heatmap, we scaled the absolute cell type prediction probabilities as z-scores across human cell types.

### Ligand receptor analysis

LR pair analysis was performed using LR analysis framework (LIANA) [82]. The human DRG non-neuronal and neuronal atlases were combined. We then ran LIANA using default settings using their human consensus database, and we filtered LR interactions with a cellphonedb p-value < 0.05 and an aggregated rank of <0.005. Both the ligand and receptor genes needed to be expressed in at least 5% of cells across the cell types used for the analysis. For each ligand-receptor interactions, the top 5 cell type pairs ranked by the aggregated rank generated by LIANA were used for visualization.

### Gene ontology analysis

We used topGO (version 2.56.0) in R (https://bioconductor.org/packages/topGO/) to perform Gene Ontology (GO) analysis. Genes of significant ligands and receptors from the ligand receptor analysis identified in the cell populations of interest were used as the input gene list. For comparison, the background gene list included all genes with average expression > 0.2 across the same populations of interest. R package org.Hs.eg.db (v3.19.1) was used as the genome-wide annotation database for human. Genes were annotated for their biological process and associated GO terms. Enrichment is defined as the number of annotated genes observed in the input list divided by the number of annotated genes expected from the background list. GO terms with more than 50 associated genes from the background list, more than 5 significant genes from the input list, and with false discovery rate of <0.05 were reported.

### In vivo electrophysiological recordings of human peripheral sensory fibers

Single-unit axonal recordings (microneurography) were obtained from the posterior antebrachial cutaneous, radial, or superficial peroneal nerve in awake, healthy participants. Participants were seated in a chair with legs or arms extended and supported using vacuum pillows—with the hand pronated for upper-limb recordings. Care was taken to ensure that each participant was comfortably seated and acclimated to the room temperature (22 °C) before starting the experiment. If participants reported feeling cold, a blanket was provided, leaving only the test region exposed.

Neural activity was sampled at 20 kHz and recorded using the ADInstruments data acquisition system (LabChart software version 8.1.24 and PowerLab 16/35 hardware, PL3516/P, Sydney, Australia). Data were either processed within LabChart or exported to Spike2 (version 10.13, Cambridge Electronic Design Ltd., Cambridge, UK) for offline analysis. Single action potentials were identified semi-automatically, with visual verification on an expanded timescale. Threshold crossing was used to distinguish action potentials from noise with a signal-to-noise ratio of at least 2:1, and spike morphology was confirmed by template matching. Recordings were discarded if multiple units were present (e.g., non-physiological interspike intervals or firing rates) or if spike amplitudes were not distinct from the noise, preventing secure action potential identification. The neural response was always spatially locked—that is, evoked (or modulated) only when the specific area of skin (the receptive field) was stimulated. Repeat trials for each stimulus were conducted to ensure reproducibility.

### Electrode placement and recording procedure

Under real-time ultrasound guidance (LOGIQ P9, GE Healthcare, Chicago, IL, USA), the target nerve was impaled with an insulated tungsten recording electrode (FHC, Inc., Bowdoin, ME, USA). Adjacent to that, an uninsulated reference electrode was inserted just under the skin. A high-impedance pre-amplifier (MLT185 Headstage, ADInstruments) was attached to the skin near the recording electrode and used together with a low-noise, high-gain amplifier (FE185 Neuro Amp EX, ADInstruments). Once the electrode tip was intrafascicular, single low-threshold mechanoreceptors (LTMRs) were searched for by soft brush stroking in the fascicular innervation zone while making fine electrode adjustments.

All recorded afferents were mechanically responsive and classified into subtypes based on established criteria (Fig. S6) [20, 56, 57]. Conduction velocity was assessed in a pooled dataset comprising previously reported units [20, 57] together with additional units collected for the present study under identical inclusion criteria and recording/stimulation protocols. Conduction velocity was estimated from response latencies to either surface electrical stimulation (FE180 Stimulus Isolator, ADInstruments) or rapid mechanical tapping using an electronic filament (Physiology Section, Department of Integrative Medical Biology, Umeå University, Sweden) applied at the receptive field. Electrically and mechanically evoked spikes were compared on an expanded timescale to confirm that they originated from the same unit.

### Mechanical stimulation of HF-LTMRs

To assess A-LTMR-hair follicles (A-LTMR[HF]) responses, individual hairs within the receptive field were deflected manually using fine forceps under magnification [58]. Air-puff stimuli were delivered using a syringe held at a short distance from the skin surface, producing a brief, localized puff of air directed onto the receptive field. The receptive field of the recorded afferent was then shaved, after which it was mechanically prodded to confirm that the recording remained stable, and responses to air puffs were compared before and after shaving. Brush stimuli were delivered using a custom-built robotic tactile stimulator (Dancer Design, St. Helens, UK), mounted on a microscope stand, which delivered high-precision rotary brush strokes at a calibrated normal force of 0.4 N [83]. Single brush strokes were applied in proximal-distal and distal-proximal directions to a 10 cm area of skin centered on the receptive field of the recorded afferent. Three velocities (1, 3, and 10 cm/s) were tested in random order. A recorded afferent was considered directionally tuned if it met the following criteria: *(1)* one brushing direction evoked a mean firing frequency across repeated trials at least 20% higher than in the opposite direction; *(2)* this difference was consistent across all tested brush velocities; and *(3)* the same preferred direction was maintained within the unit across velocities.

### Xenium *in situ* gene expression

Fresh frozen lumbar DRGs from one organ donor was gradually embedded in OCT by adding small volumes of OCT on dry ice to prevent thawing of the sample. The tissues were cryosectioned at 10 µm onto Xenium slides, then stored at –80 °C until use. Sections were fixed in 3.7% formaldehyde for 30 min at room temperature, washed in PBS, permeabilized in 1% SDS for 2 min, washed again, and incubated in 70% methanol on ice for 60 min before a final PBS wash. Following fixation and permeabilization, slides were assembled in Xenium cassettes and processed using the Xenium *In Situ* Gene Expression User Guide (10x Genomics, CG000582 Rev H) with a custom-designed 479-gene panel. The workflow included overnight probe hybridization, ligation of circularizable probes, and enzymatic amplification of gene-specific barcodes. After autofluorescence quenching and DAPI nuclear staining, samples were imaged on the Xenium Analyzer through iterative fluorescent probe binding and high-resolution imaging cycles, and transcript identities were decoded and visualized using Xenium Explorer software. Tissue preparation and fixation protocols followed the 10x Genomics demonstrated protocols for fresh-frozen tissue (CG000579, CG000581), and the full manufacturer’s workflow was adhered to in order to ensure reproducibility and data quality. Custom panel design followed 10x Genomics specifications, enabling targeted, high-plex spatial transcriptomic profiling of DRG sections.

Segmentation of lumbar DRGs from organ donors was performed using the Xenium Ranger 3.0 pipeline. Because the initial pipeline could not accurately segment large cells such as neurons, manual segmentation was carried out using the boundary staining (18S) provided in the Xenium cell segmentation kit. Neurons were delineated in Label Studio (Tkachenko et al., 2020–2025), and the re-segmented images were subsequently reprocessed with Xenium Ranger 3.0. These corrected labels were then used for downstream analysis. In contrast, segmentation of cervical and thoracic DRGs was performed using Xenium Ranger 4.0, which incorporates improved algorithms capable of segmenting larger cells, eliminating the need for manual labeling. Correctly segmented human DRG Xenium samples were integrated in Seurat (v5.0.2, R). Label transfer from a single-cell RNA-seq atlas was applied to the Xenium object using the labeltransfer function to assign neuronal subtypes within the integrated dataset. Xenium data were then visualized and exported using Xenium Explorer for downstream analyses and image generation. Neuronal diameter was estimated from the cell area feature in Xenium Explorer.

### MERSCOPE spatial transcriptomics sample preparation and imaging

One L4 DRG (male donor, 34 years; purchased from AnaBios) was mounted in optimal cutting temperature cryomount (HistoLab AB) and processed for spatial transcriptomics using the Vizgen user guide protocol for non-resistant tissue as described before [10]. In short, the tissue was sectioned with a cryostat (NX70, Thermo Fisher Scientific) in a 10 µm section and mounted on a MERSCOPE slide (Vizgen) followed by a 5 min incubation at –20 °C. Afterwards, the tissue was fixed in 5 mL 4% PFA for 15 min at room temperature and washed three times in 1X PBS for 5 minutes each. The tissue was then permeabilized in 5 mL 70% Ethanol overnight at 4 °C following a 2 minutes wash in sample prep wash buffer (Vizgen) and 30 min incubation in formamide buffer (Vizgen) at 37 °C. For probe hybridization 50 µl of a custom 300 gene probe panel (Table S4) was added to the section, covered with parafilm, and incubated at 37 °C for 48 h. Sections were washed twice in formamide buffer at 47 °C for 30 minutes and once in sample prep wash buffer for 2 min at room temperature. The tissue section was then covered in 4.9 mL gel embedding solution (5 mL gel embedding premix (Vizgen), 25 µl 10% APS, 2.5 µl TEMED) for 1 min. Solution was discarded and 50 µl Gel embedding solution was added to the section and covered with a GelSlick coated coverslip followed by a 90 min incubation at room temperature. Afterward, the coverslip was removed and the tissue was cleared in clearing solution (5 mL clearing premix (Vizgen), 50 µl Proteinase K (Sigma)) at 37 °C for 24 h. Before imaging, the section was washed twice with sample prep wash buffer and stained with DAPI and PolyT staining reagent (Vizgen) for 15 min at room temperature, washed in formamide wash buffer for 10 min, sample prep wash buffer for 5 min and immediately proceeded for imaging in the Vizgen MERSCOPE instrument following the MERSCOPE Instrument User Guide (91600001 RevG) and using the MERSCOPE 300 gene imaging kit (10400005).

### Segmentation of MERSCOPE data

For segmentation of the MERSCOPE data, the Vizgen Post Processing tool (VPT) and Cellpose2 [84] were used, following a Vizgen segmentation guide https://vizgen.github.io/vizgen-postprocessing/analysis_vignettes/segmentation_heart_dataset_cellpose2.html) in Python 3.11. In short, 15 images of DRG sections were exported and semi-manually segmented in Cellpose2. A custom model was trained based on the manual segmentation and the full dataset was resegmented with the following settings: nuclear_channel = DAPI, entity_fill_channel = PolyT, cell diameter (pixels) = 111.58, flow_threshold = 0.9, cellprob_theshold =–5.0, stitch_threshold = 0.0. The newly generated metadata files were used to update the vzg file, uploaded in the Vizgen Visualizer software, and used for visualization of the spatial transcriptomics data and exporting example images. The MERSCOPE data were then anchored to the human integrated atlas using the Seurat label transfer method.

### RNAscope in-situ hybridization

RNAscope in situ hybridization (multiplex v2) was performed according to the manufacturer’s instructions (Advanced Cell Diagnostics, ACD) along with using the 4-plex Ancillary Kit for the Multiplex Fluorescent v2 assay, and as previously described (10.1097/j.pain.0000000000001973). The probes used were RXFP1 (Channel 1, 422821), TRPV1 (Channel 2, 415381), CACNA1H (Channel 1, 528311), F10 (Channel 2, 1333641), TAC3(Channel 3, 507301), P2RY12 (Channel 4, 450391, NTRK2 (Channel 1, 402631 CALCA (Channel 2, 605551), PVALB (Channel 3, 42281), PCDH8 (Channel 1, 1218161), EPHA3 (Channel2, 1072211), FOXP2 (Channel 2, 407261), Positive control probes were included to confirm RNA integrity, and a negative control probe was used to assess background labeling. The slides were imaged on Olympus FV4000 confocal microscope at 20X magnification. The image acquisition was set based on the guidelines provided by Olympus with laser power ≤ 5%.

## DATA AND CODE AVAILABILITY

The processed sequencing data will be made publicly available on dbGAP and SPARC. Processed data will be made publicly available on the web resource. Code will be made publicly available on GitHub. All other data are either included in the supplementary tables or will be made available upon reasonable request.

## ACKNOWLEDGEMENTS

We would like to thank the patients, the donors, the donors’ families and the transplant centers, without whom much of this study would not be possible. We would also like to thank members of the Renthal, Woolf, Price, Gereau, Luo, Ernfors, and Olausson labs as well as E. Williams, H. Yong, L. He, K. Gentry, A. Salman, C. Smith, A. Shuster, S. Tripathy, and J. Gillis for helpful feedback throughout the study.

We thank the NIH HEAL PRECISION Human Pain Network for its support and for fostering a highly collaborative research environment. This work was supported by the National Institute of Neurological Disorders and Stroke [U19NS130617 (W.R. and C.J.W.), R01NS119476 (W.R.), U19NS130608 (T.J.P.), 1U19NS135528 (W.L.), and U19NS130607 (R.W.G)]. D.S.G was supported by the National Institute of General Medical Sciences of the National Institutes of Health under award number T32GM007592. W.L and single-soma RNA-seq data generation were partially supported by an Eli Lily LRAP fund. W.R. receives support from the Burroughs Wellcome Fund, Rita Allen Foundation, National Institute of Drug Abuse (DP1DA054343), National Eye Institute (U01EY034709), BWH Women’s Brain Initiative, BWH Neurotechnology Studio, and MGB Gene and Cell Therapy Institute.

## SUPPLEMENTARY TABLES

**Table S1: Donor information**

**Table S2: Neuronal marker genes generated using Seurat’s FindAllMakers function**

**Table S3: Ligand receptor interactions between neurons and non-neurons**

**Table S4: Non neuronal marker genes generated using Seurat’s FindAllMakers function**

**Table S5: Xenium and MERSCOPE probes**

**Table S6: Summary of Beta-regression modelling**

**Table S7: Spinal region-specific genes**

**Table S8: Subtype- and spinal region-specific marker genes**

**Table S9: Subtype- and sex-specific marker genes**

**Table S10: Subtype- and age-specific marker genes**

**Table S11: Human-specific marker genes for DRG neuronal subtypes**

**Table S12: Nomenclature scheme across studies**

**Table S13: Marker genes used for cell type annotation**

---

**Supplementary Fig 1.**
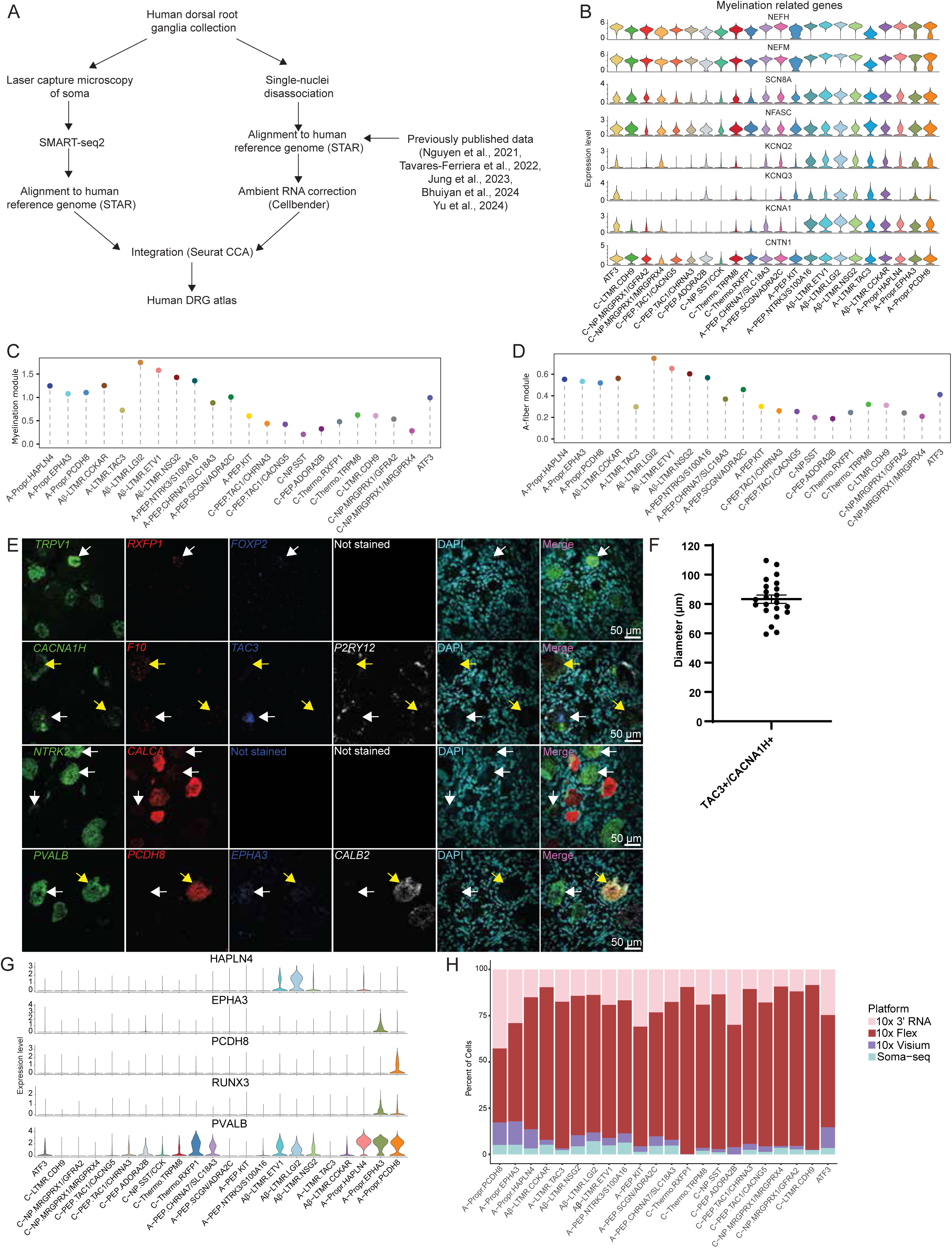
Overview of the human DRG atlas generation. (A) Workflow displays an overview of how the human DRG atlas was constructed. DRG samples were collected across the NIH PRECISION Human Pain Network. Neuronal somata were isolated via laser capture microdissection (LCM) and profiled with SMART-seq2, while single-nucleus dissociation was used for 10X Genomics-based single-nucleus RNA-seq. Both modalities were aligned to the human genome, with ambient RNA correction (CellBender) applied to 10x Genomics based data. New and previously published datasets were integrated using Seurat CCA. (B) Violin plots display log-normalized gene expression levels of canonical myelination marker genes across all identified neuronal subtypes in the human DRG atlas. (C) Dot plot displays gene module score for myelination. Gene module score was calculated using canonical myelination marker genes presented in S1B. (D) Dot plot displays gene expression profiles across human neuronal subtypes of hdWGCNA mouse module “blue” (120 co-expressed genes). These genes are strongly associated with fast conducting neuronal subtypes in mouse [10] and were used to corroborate myelination of the human neurons. (E) Representative sections display *in situ* hybridization results of *RXFP1*, *CACNA1H*, and *NTRK2* (green); *TRPV1*, *F10*, *ADAM12*, and *PCDH8* (red); *SCN10A*, *TAC3*, *CALCA*, and *EPHA3* (blue); *CHRNA7*, *P2RY12*, and *CALB2* (white); and *DAPI* (cyan). Scale bar is 50 μm across all sections. (F) Scatter plot displays the mean (±SEM) neuronal diameter of cells co-expressing *CACNA1H* and *TAC3*. Quantification of diameters of neurons were measured based on co-expression of *CACNA1H* and *TAC3* positive neurons. (G) Violin plots display the log normalized gene expression profiles of *HAPLN4, EPHA3, PCDH8, RUNX3* and *PVALB* in the human DRG neuronal atlas. (H) Bar plot displays the proportional contribution of each sequencing platform for each neuronal subtype. Bars are colored by sequencing platform.

**Supplementary Fig 2.**
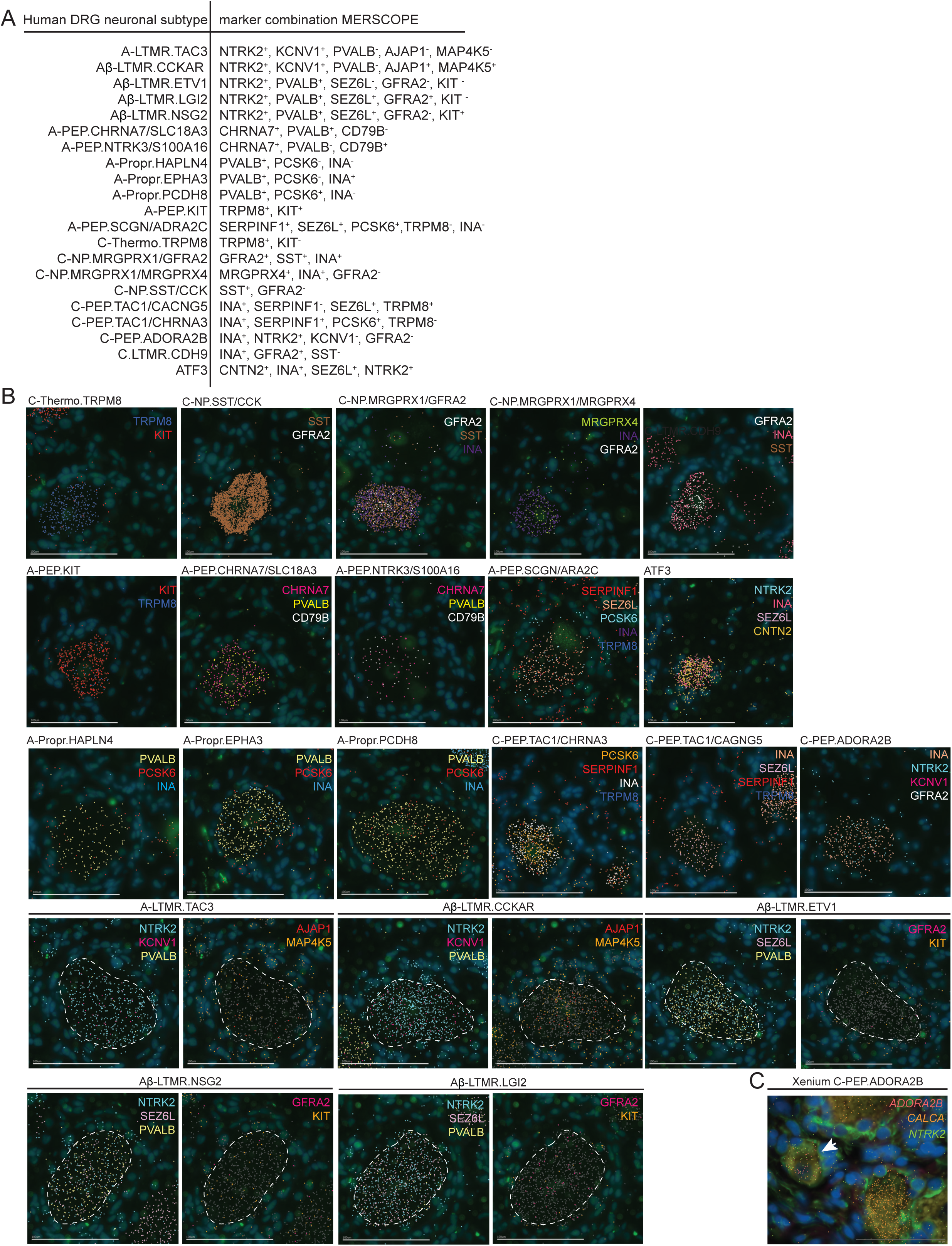
Spatial transcriptomics analysis of DRG neurons using MERSCOPE. (A) Table displays marker combinations used to identify the different human neuronal subtypes. (B) Sections display representative images of MERSCOPE transcripts used to identify neuron subpopulations. Scale bar: 100 µm. (C) Section displays representative image of Xenium transcripts used to identify C-PEP.ADORA2B. Scale bar: 50 µm.

**Supplementary Fig 3.**
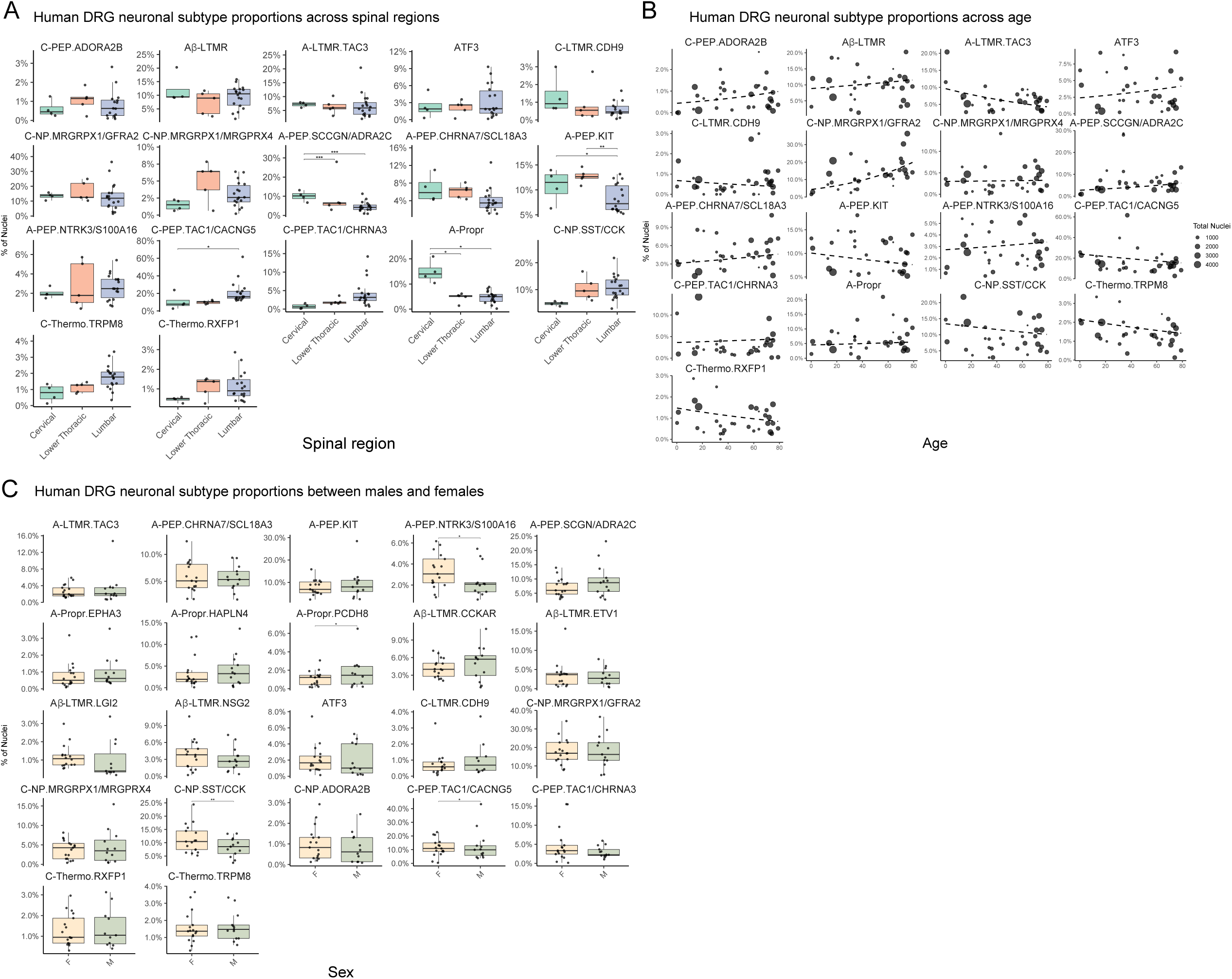
Modeling donor-level cell type proportions reveals covariate effects of age, sex, and spinal region. (A) Box plots display the proportion of neuron subtypes across three spinal levels: cervical, lower thoracic, and lumbar. Significance was calculated using a post-hoc test applied to a beta-regression model (Table S6). n.s: non-significant, *: p-value < 0.05, **: p-value<0.01, ***: p-value<0.005 (B) Scatter plots display the relationship between neuronal subtype proportions and donor age, point size displays number of cells/nuclei included in model. The dashed regression lines generated from a beta-regression model (Table S6). Significance was calculated using a post-hoc test applied to the beta-regression model. n.s: non-significant, *: p-value < 0.05, **: p-value<0.01, ***: p-value<0.005 (C) Box plots display the proportion of neuronal subtypes in female (F) and male (M) donors. Significance was calculated using a post-hoc test applied to a beta-regression model (Table S6). ns: non-significant, *: p-value < 0.05, **: p-value<0.01, ***: p-value<0.005

**Supplemental Fig. 4.**
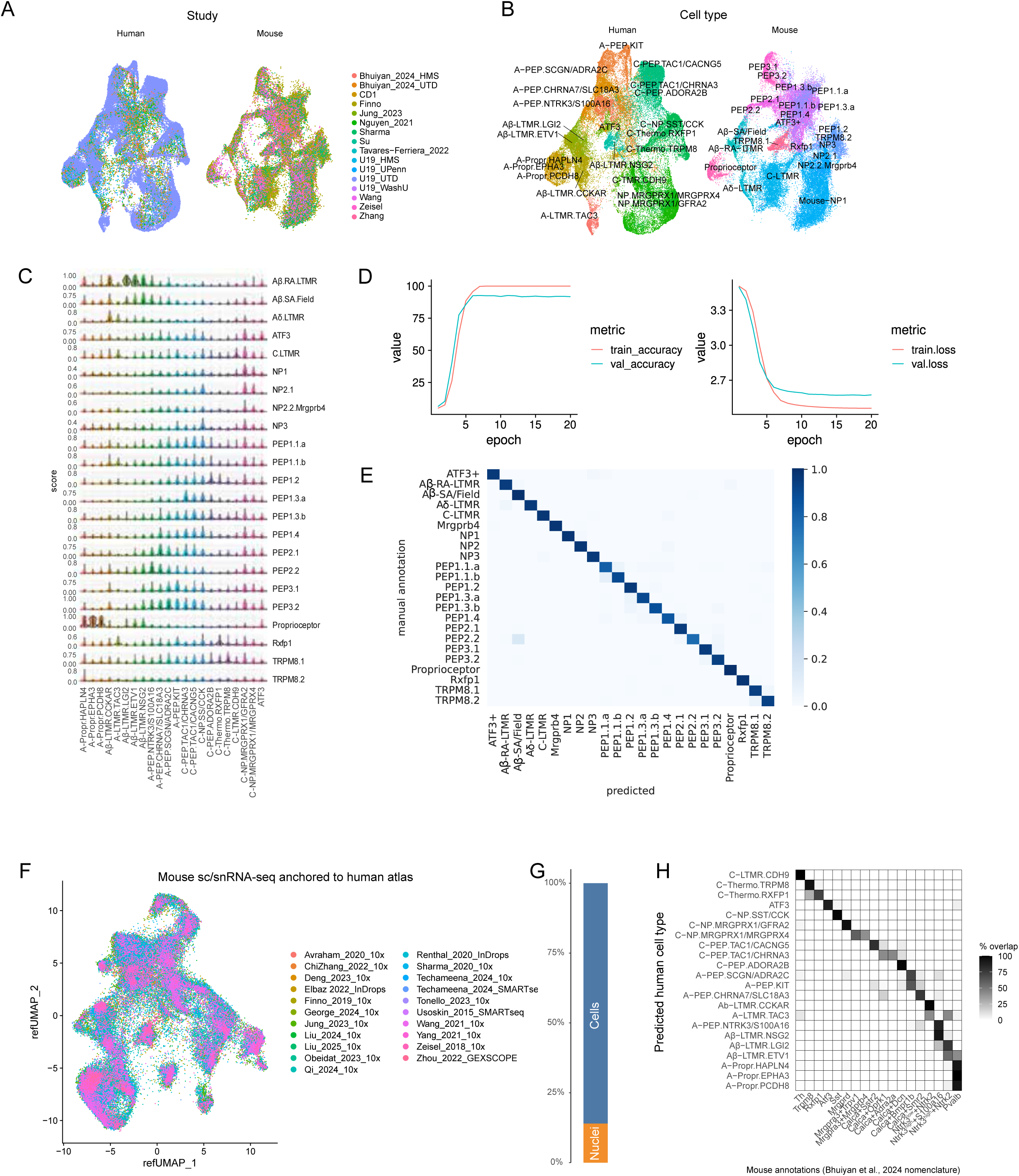
Cross-species clustering and classifier metrics. (A) UMAP displays CCA integrated mouse scRNA-seq and human data split by species and colored by data origin. (B) UMAP displays CCA integrated mouse scRNA-seq and human data split by species and colored by cell type. (C) Violin plots display the cell type prediction probability scores from the VAE model between human (columns) and mouse (rows) DRG neuron types. (D) Learning curve plot displays training metrics for the VAE model. Left: Plot displays training and validation accuracy across 20 epochs. Training accuracy increases rapidly during the first few epochs and plateaus near 100%, while validation accuracy follows a similar trajectory and stabilizes slightly below the training curve. Right: Plot shows training and validation loss across 20 epochs. Both training and validation loss decrease sharply during the initial epochs and then level off, indicating convergence of the model without substantial overfitting. (E) Heatmap displays VAE model performance on 20% of the held-out test data, comparing predicted neuronal subtypes (x-axis) to manual annotations (y-axis). Each cell represents the percentage of nuclei assigned to a given predicted class. Strong diagonal values indicate high concordance between predicted and manually annotated labels, demonstrating accurate subtype classification. (F) UMAP displays an integrated mouse atlas of 21 previously published single-cell and single-nucleus RNA-sequencing studies. Cells/nuclei are colored by study of origin (“study_sequencing platform”), and a total of 103,188 cells/nuclei are displayed. (G) Bar plot displays the proportion of sc/snRNA-seq data (panel F) that are sequenced cells versus sequenced nuclei. (H) Heatmap displays the percentage of predicted mouse cell types (x-axis) for each human cell type (y-axis). The human atlas was anchored to the mouse sc/snRNA-seq data using the Seurat label transfer approach (see methods). Mouse cell types were labeled based on the Bhuiyan et al., 2024 nomenclature scheme (Figure 6C; [9]). Cells/nuclei with a prediction score > 0.6 are displayed.

**Figure S5.**
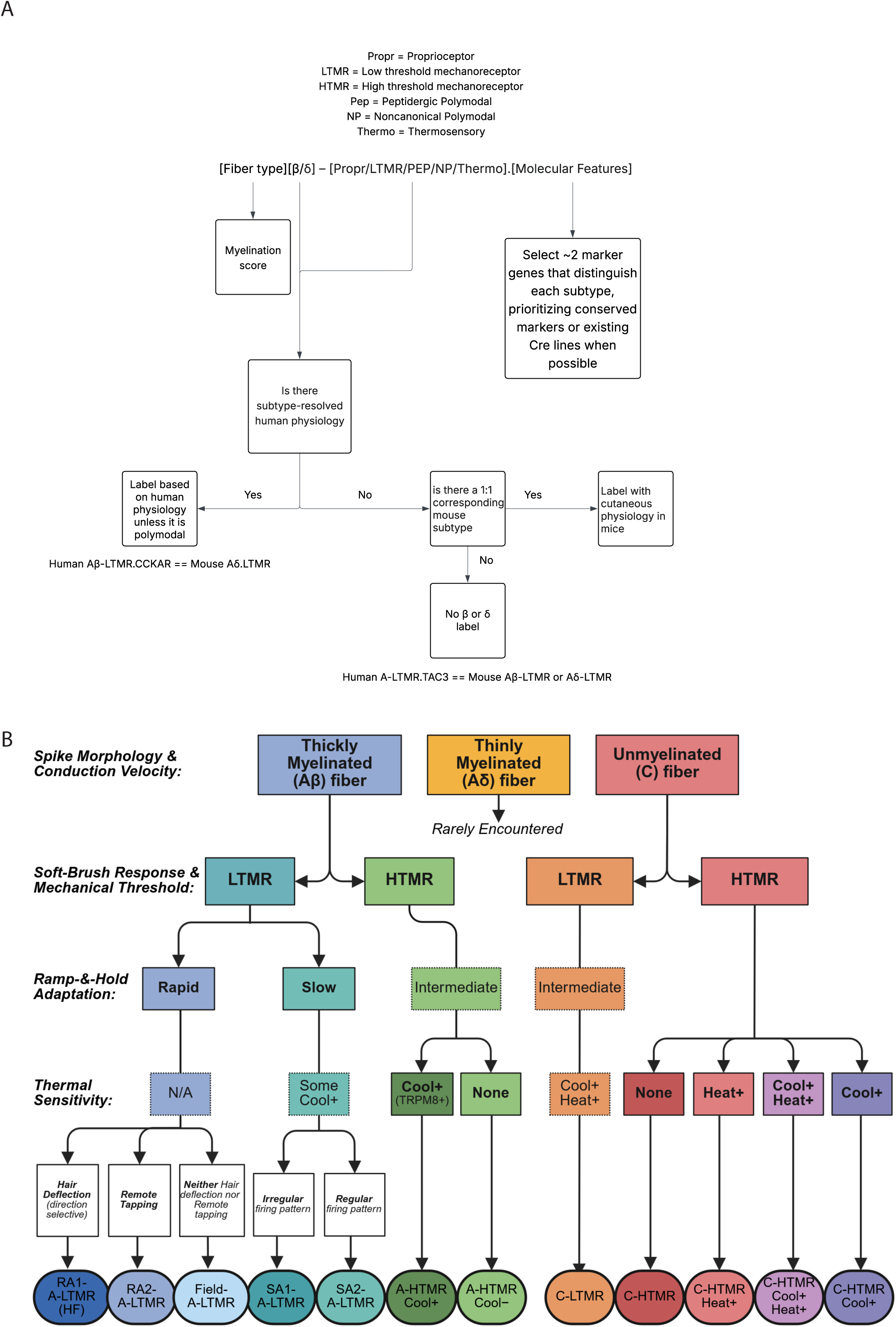
Nomenclature workflow. (A) Flowchart displays the framework used to assign fiber type and functional subtype labels to human DRG neuronal clusters. Nomenclature is structured as [Fiber type][β/δ]-[Physiology].[Marker genes]. <u>Fiber type</u>: Myelination scores are first used to infer fiber type (A- vs C-fiber). <u>β/δ and Physiology</u>: When subtype-resolved human physiology was available, labels were based on known human functional properties unless the subtype was polymodal. For subtypes lacking human physiological data, labels were transferred from one-to-one corresponding mouse subtypes when present. If no corresponding mouse subtype was identified, no β or δ label was assigned. <u>Marker genes</u>: Approximately two marker genes distinguishing each subtype were selected, prioritizing conserved markers or existing Cre lines when possible. The ATF3 cell type was excluded from this workflow. This cell type has been identified in the naïve state in at least six species; however, it expresses molecular features of any DRG neuron after peripheral nerve injury, regardless of fiber type. As ATF3 may be neurons in an injury state or a naïve cell type, we chose to not assign ATF3 with a fiber type. (B) Workflow displays how physiologically distinct cell types are assigned. Human mechanosensitive afferents recorded using single-fiber microneurography are classified by spike shape (A or C), conduction velocity (Aβ, Aδ, or C), and response properties, including sensitivity to soft brushing (present/absent), von Frey threshold (low/high), and adaptation to sustained indentation (rapid/intermediate/slow), and are further subclassed based on thermal responsiveness (cooling/heating). Aβ afferents include low-threshold mechanoreceptors (LTMRs: RA1/HF, RA2, Field, SA1, SA2) and high-threshold mechanoreceptors (HTMRs: cool* or cool̅). Aδ fibers are rarely encountered. Unmyelinated C fibers comprise LTMRs and HTMRs with distinct thermal sensitivities (heat*, cool*, both cool* and heat*, or none). Features in bold denote criteria used in unit classification, while plain text represents additional observed properties.

## REFERENCES

1. Kupari J, Ernfors P. Molecular taxonomy of nociceptors and pruriceptors. PAIN. 2022;:10.1097/j.pain.0000000000002831. 10.1097/j.pain.0000000000002831.

2. Qi L, Iskols M, Shi D, Reddy P, Walker C, Lezgiyeva K, et al. A mouse DRG genetic toolkit reveals morphological and physiological diversity of somatosensory neuron subtypes. Cell. 2024. 10.1016/j.cell.2024.02.006.

3. Yang L, Xu M, Bhuiyan SA, Li J, Zhao J, Cohrs RJ, et al. Human and mouse trigeminal ganglia cell atlas implicates multiple cell types in migraine. Neuron. 2022. 10.1016/j.neuron.2022.03.003.

4. Dydyk AM, Conermann T. Chronic Pain. In: StatPearls. Treasure Island (FL): StatPearls Publishing; 2024.

5. Adriaensen H, Gybels J, Handwerker HO, Van Hees J. Response properties of thin myelinated (A-delta) fibers in human skin nerves. J Neurophysiol. 1983;49:111–22. 10.1152/jn.1983.49.1.111.

6. Usoskin D, Furlan A, Islam S, Abdo H, Lönnerberg P, Lou D, et al. Unbiased classification of sensory neuron types by large-scale single-cell RNA sequencing. Nat Neurosci. 2015;18:145–53. 10.1038/nn.3881.

7. Zeisel A, Hochgerner H, Lönnerberg P, Johnsson A, Memic F, van der Zwan J, et al. Molecular Architecture of the Mouse Nervous System. Cell. 2018;174:999–1014.e22. 10.1016/j.cell.2018.06.021.

8. Kupari J, Usoskin D, Parisien M, Lou D, Hu Y, Fatt M, et al. Single cell transcriptomics of primate sensory neurons identifies cell types associated with chronic pain. Nat Commun. 2021;12:1510. 10.1038/s41467-021-21725-z.

9. Bhuiyan SA, Xu M, Yang L, Semizoglou E, Bhatia P, Pantaleo KI, et al. Harmonized cross-species cell atlases of trigeminal and dorsal root ganglia. Sci Adv. 2024;10:eadj9173. 10.1126/sciadv.adj9173.

10. Krauter D, Kupari J, Usoskin D, Su J, Hu Y, Zhang M-D, et al. Spatial organization, chromatin accessibility and gene-regulatory programs defining mouse sensory neurons. Commun Biol. 2025;8:1–19. 10.1038/s42003-025-08315-1.

11. Chesler AT, Szczot M, Bharucha-Goebel D, Čeko M, Donkervoort S, Laubacher C, et al. The Role of PIEZO2 in Human Mechanosensation. N Engl J Med. 2016;375:1355–64. 10.1056/NEJMoa1602812.

12. Renthal W, Tochitsky I, Yang L, Cheng Y-C, Li E, Kawaguchi R, et al. Transcriptional Reprogramming of Distinct Peripheral Sensory Neuron Subtypes after Axonal Injury. Neuron. 2020;108:128–144.e9. 10.1016/j.neuron.2020.07.026.

13. Ghitani N, von Buchholtz LJ, MacDonald DI, Falgairolle M, Nguyen MQ, Licholai JA, et al. A distributed coding logic for thermosensation and inflammatory pain. Nature. 2025;642:1016–23. 10.1038/s41586-025-08875-6.

14. MacDonald DI, Jayabalan M, Seaman JT, Balaji R, Nickolls AR, Chesler AT. Pain persists in mice lacking both Substance P and CGRPa signaling. eLife. 2025;13:RP93754. 10.7554/eLife.93754.

15. Renthal W, Chamessian A, Curatolo M, Davidson S, Burton M, Dib-Hajj S, et al. Human cells and networks of pain: Transforming pain target identification and therapeutic development. Neuron. 2021;109:1426–9. 10.1016/j.neuron.2021.04.005.

16. Copits BA, Curatolo M, Dougherty PM, Gereau IV RW, Luo W, Martone M, et al. Human pain neuroscience and the next generation of pain therapeutics. Neuron. 2025;113:1304–6. 10.1016/j.neuron.2025.04.005.

17. Nguyen MQ, von Buchholtz LJ, Reker AN, Ryba NJ, Davidson S. Single-nucleus transcriptomic analysis of human dorsal root ganglion neurons. eLife. 2021;10:e71752. 10.7554/eLife.71752.

18. Tavares-Ferreira D, Shiers S, Ray PR, Wangzhou A, Jeevakumar V, Sankaranarayanan I, et al. Spatial transcriptomics of dorsal root ganglia identifies molecular signatures of human nociceptors. Sci Transl Med. 2022;14:eabj8186. 10.1126/scitranslmed.abj8186.

19. Jung M, Dourado M, Maksymetz J, Jacobson A, Laufer BI, Baca M, et al. Cross-species transcriptomic atlas of dorsal root ganglia reveals species-specific programs for sensory function. Nat Commun. 2023;14:366. 10.1038/s41467-023-36014-0.

20. Yu H, Nagi SS, Usoskin D, Hu Y, Kupari J, Bouchatta O, et al. Leveraging deep single-soma RNA sequencing to explore the neural basis of human somatosensation. Nat Neurosci. 2024;27:2326–40. 10.1038/s41593-024-01794-1.

21. Avraham O, Chamessian A, Feng R, Yang L, Halevi AE, Moore AM, et al. Profiling the molecular signature of satellite glial cells at the single cell level reveals high similarities between rodents and humans. Pain. 2022;163:2348–64. 10.1097/j.pain.0000000000002628.

22. Wu Z, Wang Y, Chen W, Sun H, Chen X, Li X, et al. Peripheral nervous system microglia-like cells regulate neuronal soma size throughout evolution. Cell. 2025;188:2159–2174.e15. 10.1016/j.cell.2025.02.007.

23. Avraham O, Deng P-Y, Jones S, Kuruvilla R, Semenkovich CF, Klyachko VA, et al. Satellite glial cells promote regenerative growth in sensory neurons. Nat Commun. 2020;11:4891. 10.1038/s41467-020-18642-y.

24. Zhang D, Wei Y, Liu J, Yang Y, Ou M, Chen Y, et al. Single-nucleus transcriptomic analysis reveals divergence of glial cells in peripheral somatosensory system between human and mouse. 2022;:2022.02.15.480622. 10.1101/2022.02.15.480622.

25. Carr MJ, Toma JS, Johnston APW, Steadman PE, Yuzwa SA, Mahmud N, et al. Mesenchymal Precursor Cells in Adult Nerves Contribute to Mammalian Tissue Repair and Regeneration. Cell Stem Cell. 2019;24:240–256.e9. 10.1016/j.stem.2018.10.024.

26. Lovatt D, Tamburino A, Krasowska-Zoladek A, Sanoja R, Li L, Peterson V, et al. scRNA-seq generates a molecular map of emerging cell subtypes after sciatic nerve injury in rats. Commun Biol. 2022;5:1105. 10.1038/s42003-022-03970-0.

27. Heming M, Börsch A-L, Wolbert J, Thomas C, Mausberg AK, Szepanowski F, et al. Multi-omic identification of perineurial hyperplasia and lipid-associated nerve macrophages in human polyneuropathies. Nat Commun. 2025;16:7872. 10.1038/s41467-025-62964-8.

28. Wang PL, Yim AKY, Kim K-W, Avey D, Czepielewski RS, Colonna M, et al. Peripheral nerve resident macrophages share tissue-specific programming and features of activated microglia. Nat Commun. 2020;11:2552. 10.1038/s41467-020-16355-w.

29. Hakim S, Jain A, Adamson SS, Petrova V, Indajang J, Kim HW, et al. Macrophages protect against sensory axon loss in peripheral neuropathy. Nature. 2025;640:212–20. 10.1038/s41586-024-08535-1.

30. Ji R-R, Chamessian A, Zhang Y-Q. Pain regulation by non-neuronal cells and inflammation. Science. 2016;354:572–7. 10.1126/science.aaf8924.

31. Lund H, Hunt MA, Kurtović Z, Sandor K, Kägy PB, Fereydouni N, et al. CD163+ macrophages monitor enhanced permeability at the blood-dorsal root ganglion barrier. J Exp Med. 2024;221:e20230675. 10.1084/jem.20230675.

32. Niimi K, Nakae J, Kubota Y, Inagaki S, Furuyama T. Macrophages play a crucial role in vascular smooth muscle cell coverage. Development. 2024;151:dev203080. 10.1242/dev.203080.

33. Fukuda S, Broxmeyer HE, Pelus LM. Flt3 ligand and the Flt3 receptor regulate hematopoietic cell migration by modulating the SDF-1alpha(CXCL12)/CXCR4 axis. Blood. 2005;105:3117–26. 10.1182/blood-2004-04-1440.

34. Wang J-G, Strong JA, Xie W, Zhang J-M. Local Inflammation in Rat Dorsal Root Ganglion Alters Excitability and Ion Currents in Small Diameter Sensory Neurons. Anesthesiology. 2007;107:322–32. 10.1097/01.anes.0000270761.99469.a7.

35. Zigmond RE, Echevarria FD. Macrophage biology in the peripheral nervous system after injury. Prog Neurobiol. 2019;173:102–21. 10.1016/j.pneurobio.2018.12.001.

36. Jain A, Gyori BM, Hakim S, Jain A, Sun L, Petrova V, et al. Nociceptor-immune interactomes reveal insult-specific immune signatures of pain. Nat Immunol. 2024;25:1296–305. 10.1038/s41590-024-01857-2.

37. Reza JN, Gavazzi I, Cohen J. Neuropilin-1 Is Expressed on Adult Mammalian Dorsal Root Ganglion Neurons and Mediates Semaphorin3a/Collapsin-1-Induced Growth Cone Collapse by Small Diameter Sensory Afferents. Mol Cell Neurosci. 1999;14:317–26. 10.1006/mcne.1999.0786.

38. Gomez K, Duran P, Tonello R, Allen HN, Boinon L, Calderon-Rivera A, et al. Neuropilin-1 is essential for VEGFA-mediated increase of sensory neuron activity and development of pain-like behaviors. Pain. 2023;164:2696–710. 10.1097/j.pain.0000000000002970.

39. Peach CJ, Tonello R, Damo E, Gomez K, Calderon-Rivera A, Bruni R, et al. Neuropilin-1 is a co-receptor for Nerve Growth Factor-evoked pain. 2024;:2023.12.06.570398. 10.1101/2023.12.06.570398.

40. Wu H, Petitpré C, Fontanet P, Sharma A, Bellardita C, Quadros RM, et al. Distinct subtypes of proprioceptive dorsal root ganglion neurons regulate adaptive proprioception in mice. Nat Commun. 2021;12:1026. 10.1038/s41467-021-21173-9.

41. Kiasalari Z, Salehi I, Zhong Y, MCmahon SB, Michael-Titus AT, Michael GJ. Identification of perineal sensory neurons activated by innocuous heat. J Comp Neurol. 2010;518:137–62. 10.1002/cne.22187.

42. Wolfson RL, Abdelaziz A, Rankin G, Kushner S, Qi L, Mazor O, et al. DRG afferents that mediate physiologic and pathologic mechanosensation from the distal colon. Cell. 2023;186:3368–3385.e18. 10.1016/j.cell.2023.07.007.

43. Mallo M, Wellik DM, Deschamps J. Hox genes and regional patterning of the vertebrate body plan. Dev Biol. 2010;344:7–15. 10.1016/j.ydbio.2010.04.024.

44. Mallo M. Reassessing the Role of Hox Genes during Vertebrate Development and Evolution. Trends Genet. 2018;34:209–17. 10.1016/j.tig.2017.11.007.

45. Simon A, Lofthouse RA, Miti P, Giuraniuc CV, Banks RW, Bewick GS. Calcium regulation of muscle spindle mechanosensory afferent function. Exp Physiol. n/a n/a. 10.1113/EP092558.

46. Mecklenburg J, Zou Y, Wangzhou A, Garcia D, Lai Z, Tumanov AV, et al. Transcriptomic sex differences in sensory neuronal populations of mice. Sci Rep. 2020;10:15278. 10.1038/s41598-020-72285-z.

47. Shen BQ, Sankaranarayanan I, Price TJ, Tavares-Ferreira D. Sex-differences in prostaglandin signaling: a semi-systematic review and characterization of PTGDS expression in human sensory neurons. Sci Rep. 2023;13:4670. 10.1038/s41598-023-31603-x.

48. Franco-Enzástiga Ú, Inturi NN, Natarajan K, Mwirigi JM, Mazhar K, Schlachetzki JCM, et al. Epigenomic landscape of the human dorsal root ganglion: sex differences and transcriptional regulation of nociceptive genes. 2024;:2024.03.27.587047. 10.1101/2024.03.27.587047.

49. Harkins SW, Price DD, Martelli M. Effects of Age on Pain Perception: Thermonociception1. J Gerontol. 1986;41:58–63. 10.1093/geronj/41.1.58.

50. Yezierski RP. The Effects of Age on Pain Sensitivity: Preclinical Studies. Pain Med. 2012;13 suppl_2:S27–36. 10.1111/j.1526-4637.2011.01311.x.

51. Colloca L, Ludman T, Bouhassira D, Baron R, Dickenson AH, Yarnitsky D, et al. Neuropathic pain. Nat Rev Dis Primer. 2017;3:17002. 10.1038/nrdp.2017.2.

52. Kenshalo DR. Somesthetic sensitivity in young and elderly humans. J Gerontol. 1986;41:732–42. 10.1093/geronj/41.6.732.

53. Feng J, Luo J, Yang P, Du J, Kim BS, Hu H. Piezo2 channel-Merkel cell signaling modulates the conversion of touch to itch. Science. 2018;360:530–3. 10.1126/science.aar5703.

54. van Reij RRI, Hoofwijk DMN, Rutten BPF, Weinhold L, Leber M, Joosten E a. J, et al. The association between genome-wide polymorphisms and chronic postoperative pain: a prospective observational study. Anaesthesia. 2020;75:e111–20. 10.1111/anae.14832.

55. Bonomo RR, Cook TM, Gavini CK, White CR, Jones JR, Bovo E, et al. Fecal transplantation and butyrate improve neuropathic pain, modify immune cell profile, and gene expression in the PNS of obese mice. Proc Natl Acad Sci. 2020;117:26482–93. 10.1073/pnas.2006065117.

56. Vallbo AB, Olausson H, Wessberg J, Kakuda N. Receptive field characteristics of tactile units with myelinated afferents in hairy skin of human subjects. J Physiol. 1995;483 (Pt 3) Pt 3:783–95. 10.1113/jphysiol.1995.sp020622.

57. Nagi SS, Marshall AG, Makdani A, Jarocka E, Liljencrantz J, Ridderström M, et al. An ultrafast system for signaling mechanical pain in human skin. Sci Adv. 2019;5:eaaw1297. 10.1126/sciadv.aaw1297.

58. Moore W, Nikesjö J, Bouchatta O, Makdani AD, Hakizimana P, Rousson M, et al. Robust coupling between the C-tactile afferent and the hair follicle in humans. J Physiol. 2025;603:4593–608. 10.1113/JP287706.

59. Rutlin M, Ho C-Y, Abraira VE, Cassidy C, Bai L, Woodbury CJ, et al. The Cellular and Molecular Basis of Direction Selectivity of Aδ-LTMRs. Cell. 2014;159:1640–51. 10.1016/j.cell.2014.11.038.

60. Paricio-Montesinos R, Schwaller F, Udhayachandran A, Rau F, Walcher J, Evangelista R, et al. The Sensory Coding of Warm Perception. Neuron. 2020;106:830–841.e3. 10.1016/j.neuron.2020.02.035.

61. Bautista DM, Siemens J, Glazer JM, Tsuruda PR, Basbaum AI, Stucky CL, et al. The menthol receptor TRPM8 is the principal detector of environmental cold. Nature. 2007;448:204–8. 10.1038/nature05910.

62. Dhaka A, Murray AN, Mathur J, Earley TJ, Petrus MJ, Patapoutian A. TRPM8 Is Required for Cold Sensation in Mice. Neuron. 2007;54:371–8. 10.1016/j.neuron.2007.02.024.

63. Tan C-H, McNaughton PA. The TRPM2 ion channel is required for sensitivity to warmth. Nature. 2016;536:460–3. 10.1038/nature19074.

64. Campero M, Baumann TK, Bostock H, Ochoa JL. Human cutaneous C fibres activated by cooling, heating and menthol. J Physiol. 2009;587 Pt 23:5633–52. 10.1113/jphysiol.2009.176040.

65. Yu H, Zhao T, Liu S, Wu Q, Johnson O, Wu Z, et al. MRGPRX4 is a bile acid receptor for human cholestatic itch. eLife. 2019;8:e48431. 10.7554/eLife.48431.

66. Bouchatta O, Brodzki M, Manouze H, Carballo GB, Kindström E, de-Faria FM, et al. PIEZO2-dependent rapid pain system in humans and mice. 2023;:2023.12.01.569650. 10.1101/2023.12.01.569650.

67. Oliver KM, Florez-Paz DM, Badea TC, Mentis GZ, Menon V, de Nooij JC. Molecular correlates of muscle spindle and Golgi tendon organ afferents. Nat Commun. 2021;12:1451. 10.1038/s41467-021-21880-3.

68. Dietrich S, Company C, Song K, Lowenstein ED, Riedel L, Birchmeier C, et al. Molecular identity of proprioceptor subtypes innervating different muscle groups in mice. Nat Commun. 2022;13:6867. 10.1038/s41467-022-34589-8.

69. Richmond FJ, Abrahams VC. What are the proprioceptors of the neck? Prog Brain Res. 1979;50:245–54. 10.1016/S0079-6123(08)60825-0.

70. Handler A, Ginty DD. The mechanosensory neurons of touch and their mechanisms of activation. Nat Rev Neurosci. 2021;22:521–37. 10.1038/s41583-021-00489-x.

71. Luo W, Enomoto H, Rice FL, Milbrandt J, Ginty DD. Molecular Identification of Rapidly Adapting Mechanoreceptors and their Developmental Dependence on Ret Signaling. Neuron. 2009;64:841–56. 10.1016/j.neuron.2009.11.003.

72. Hodge RD, Bakken TE, Miller JA, Smith KA, Barkan ER, Graybuck LT, et al. Conserved cell types with divergent features in human versus mouse cortex. Nature. 2019;573:61–8. 10.1038/s41586-019-1506-7.

73. Treede R-D, Rief W, Barke A, Aziz Q, Bennett MI, Benoliel R, et al. Chronic pain as a symptom or a disease: the IASP Classification of Chronic Pain for the International Classification of Diseases (ICD-11). Pain. 2019;160:19–27. 10.1097/j.pain.0000000000001384.

74. North RY, Li Y, Ray P, Rhines LD, Tatsui CE, Rao G, et al. Electrophysiological and transcriptomic correlates of neuropathic pain in human dorsal root ganglion neurons. Brain J Neurol. 2019;142:1215–26. 10.1093/brain/awz063.

75. Sankaranarayanan I, Price T. Single nuclei RNA sequencing from human dorsal root ganglion. 2025.

76. Heaton H, Talman AM, Knights A, Imaz M, Gaffney DJ, Durbin R, et al. Souporcell: robust clustering of single-cell RNA-seq data by genotype without reference genotypes. Nat Methods. 2020;17:615–20. 10.1038/s41592-020-0820-1.

77. Rubinacci S, Hofmeister RJ, Sousa da Mota B, Delaneau O. Imputation of low-coverage sequencing data from 150,119 UK Biobank genomes. Nat Genet. 2023;55:1088–90. 10.1038/s41588-023-01438-3.

78. Koenig Z, Yohannes MT, Nkambule LL, Zhao X, Goodrich JK, Kim HA, et al. A harmonized public resource of deeply sequenced diverse human genomes. BioRxiv Prepr Serv Biol. 2024;:2023.01.23.525248. 10.1101/2023.01.23.525248.

79. Boyer K, Meriau P, Yang L, Naz H, Rosen S, Murray G, et al. Single nucleus multiomic atlas of human dorsal root ganglia reveals the contribution of non-neuronal cell types to pain. 2025;:2025.09.21.677655. 10.1101/2025.09.21.677655.

80. Yang L, Meriau P, Cavalli V, Gereau RW. Nuclear extraction using opti-prep gradient centrifugation for 10X Genomics Multiome Assay. 2025.

81. Pollock JD, Wu D-Y, Satterlee JS. Molecular neuroanatomy: a generation of progress. Trends Neurosci. 10.1016/j.tins.2013.11.001.

82. Dimitrov D, Türei D, Garrido-Rodriguez M, Burmedi PL, Nagai JS, Boys C, et al. Comparison of methods and resources for cell-cell communication inference from single-cell RNA-Seq data. Nat Commun. 2022;13:3224. 10.1038/s41467-022-30755-0.

83. Löken LS, Wessberg J, Morrison I, McGlone F, Olausson H. Coding of pleasant touch by unmyelinated afferents in humans. Nat Neurosci. 2009;12:547–8. 10.1038/nn.2312.

84. Pachitariu M, Stringer C. Cellpose 2.0: how to train your own model. Nat Methods. 2022;19:1634–41. 10.1038/s41592-022-01663-4.

